# Enhanced cell aggregation in the *Chlamydomonas reinhardtii rbo1* mutant in response to multifactorial stress combination

**DOI:** 10.1101/2025.05.16.654601

**Authors:** Lidia S. Pascual, Devasantosh Mohanty, Ranjita Sinha, Thi Thao Nguyen, Linda Rowland, Zhen Lyu, Trupti Joshi, Brian P. Mooney, Aurelio Gómez-Cadenas, Felix B. Fritschi, Sara I. Zandalinas, Ron Mittler

## Abstract

Global change factors associated with climate change and increased pollution are subjecting plants, microbes, and different ecosystems to conditions of multifactorial stress combination (MFSC). While recent studies centered on the effects of MFSC on different plants and ecosystems, much less is known about how these conditions impact unicellular organisms. Here, we report on the physiological and proteomic responses of wild type and respiratory burst oxidase homolog 1 (*rbo1*) mutant cells of *Chlamydomonas reinhardtii* to a MFSC of 5 different abiotic stresses. While several similarities were found between plant and *C. reinhardtii* responses to MFSC, our work revealed that MFSC induces aggregation in the unicellular organism *C. reinhardtii*. We further show that MFSC induced aggregation is enhanced in the *rbo1* mutant and can be triggered by H_2_O_2_. As aggregation typically leads to reduced growth, respiration and/or photosynthesis, enhanced aggregation of unicellular organisms in different ecosystems subjected to MFSC could explain part of the negative impacts of MFSC on the services they provide. Our findings shed new light on the response of unicellular organisms to MFSC and suggest that one of the main drivers leading to multicellularity during evolution was early conditions of MFSC under an oxygenated environment.

## Introduction

Anthropogenic activities are altering the environmental conditions on our planet, subjecting different unicellular and multicellular organisms to a multitude of different stressors, simultaneously or sequentially (Lesk et al., 2016; Sage, 2020; Intergovernmental Panel on Climate Change, 2023; Richardson et al., 2023). Examples for such stress conditions include elevated day and night temperatures, increased levels of water, soil, and air pollutants, and in some areas of our planet, higher salinity levels, associated with flooding, droughts, and/or different agricultural practices (Mittler, 2006; Bailey-Serres et al., 2019; Zhang et al., 2021; Rhee et al., 2024). Recent studies revealed that with the increased number of different stressors, or global change factors (GCFs), simultaneously impacting an ecosystem (*i.e.,* GCF combination; GCFc), the overall services provided by the ecosystem and its biological diversity significantly decline (Rillig et al., 2019; Rillig et al., 2021; Speißer et al., 2022). Similarly, recent studies conducted with Arabidopsis (*Arabidopsis thaliana*), tomato (*Solanum lycopersicum*), soybean (*Glycine max*), rice (*Oryza sativa*), and corn (*Zea mays*), subjected to a combination of multiple abiotic stresses (*i.e.,* multifactorial stress combination; MFSC), revealed that with the increased number of stresses simultaneously impacting a plant, plant growth, physiological activity, yield, and/or survival significantly decline (Zandalinas et al., 2021; Pascual et al., 2023; Peláez-Vico et al., 2023; Sinha et al., 2024). Taken together, the studies referenced above, reveal a new principle in biology, termed ‘the GCFc or MFSC principle’. This principle states that with the increased number, or complexity, of environmental stressors simultaneously impacting an organism or an ecosystem, the health of the organism or ecosystem will significantly (and dramatically) decline, even if the level of each stressor, when individually applied to the plant or ecosystem, is minimal (Rillig et al., 2019; Zandalinas et al., 2021; Zandalinas and Mittler, 2022).

Phenotypic, genetic, transcriptomic, proteomic, and/or mixomics analyses of Arabidopsis, soybean, and rice, subjected to MFSC suggested that pathways essential for the maintenance of reactive oxygen species (ROS), Fe, Cu, 2Fe-2S, and ascorbic acid homeostasis are crucial for plant survival under conditions of MFSC (Zandalinas et al., 2021; Peláez-Vico et al., 2023; Sinha et al., 2024). In addition, the expression level and abundance of many transcripts and proteins with unknown function were found to be elevated under conditions of MFSC (Zandalinas et al., 2021; Peláez-Vico et al., 2023; Sinha et al., 2024). These findings suggest that unique pathways, networks, and/or genes might be required for plant acclimation to MFSC.

Although the negative impacts of GCFc on the diversity and respiration of soil microorganisms was studied (Rillig et al., 2019), little is known about how a single unicellular organism will respond at the physiological, molecular, and biochemical levels to MFSC. To study the response of a unicellular organism to MFSC, and compare it to that of multicellular organisms, we chose the unicellular green algae *Chlamydomonas reinhardtii*. *C. reinhardtii* belongs to an algal unicellular family that grows in different aquatic ecosystems, soils, and even on snow, and is likely subjected in nature to multiple stressors in different combinations, including salinity, different types of pollutants, temperature extremes, and cycles of hydration and dehydration (Li et al., 2013; Liu et al., 2014; Sasso et al., 2018; Hoham and Remias, 2020). *C. reinhardtii* is also a photosynthetic eukaryotic green alga and most, if not all, the molecular and physiological data we have access to from multicellular organisms subjected to MFSC comes from higher plants (Zandalinas et al., 2021; Pascual et al., 2023; Peláez-Vico et al., 2023; Sinha et al., 2024). *C. reinhardtii* is therefore an excellent candidate for a comparison between the responses of a photosynthetic unicellular organism (*i.e., C. reinhardtii*) to that of a photosynthetic multicellular organism (*i.e.,* a higher plant) to MFSC.

## Results and Discussion

### The impact of MFSC on the growth, chlorophyll content and extracellular H_2_O_2_ levels of *C. reinhardtii*

To study the responses of *C. reinhardtii* to MFSC, we subjected wild type (WT) *C. reinhardtii* and a mutant of *C. reinhardtii*, deficient in respiratory burst oxidase homolog 1 (*rboh1*, hereafter referred to as *rbo1*; Anderson et al., 2016), to a MFSC using liquid medium conditions. The *rbo1* mutant was previously shown to have suppressed ROS production and impaired cell-to-cell ROS signaling in response to an excess light stress treatment, when grown as a lawn on agar plates (Fichman et al., 2023; Zhou et al., 2024). For MFSC, we used the following conditions: salinity (120 mM NaCl; S), the heavy metal cadmium (300 µM; Cd), excess light (700 μmol m^−2^ s^−1^; EL; a 7-fold increase over control growth conditions), acidic pH (pH 5.5; pH), and the herbicide paraquat (50 nM; PQ), applied for 24 hours (no changes in media pH were found following the stress treatments). Similar to experimental designs we used for higher plants (Zandalinas et al., 2021; Pascual et al., 2023; Peláez-Vico et al., 2023; Sinha et al., 2024), S, Cd, EL, and PQ were applied in all possible combinations, with pH added to the 4-stress combination to generate a 5-stress combination treatment.

While most stresses (except EL) applied individually (1-stress condition) had a significant effect on the growth rate of WT and the *rbo1* mutant, the impact of 2-stress combinations was significantly more severe, and the impact of 3-stress combinations reached the maximum negative effect, that was similar to the impact of 4- and 5-stress combination treatments (Figures 1A, S1). No differences were observed between the WT and the *rbo1* mutant under any of the stress treatments (Figure 1A). The impact of MFSC on the growth rate of *C. reinhardtii* displayed therefore a similar pattern to that of MFSC on the growth of flowering plants (Zandalinas et al., 2021; Pascual et al., 2023; Peláez-Vico et al., 2023; Sinha et al., 2024). Interestingly, while differences were observed between WT and the respiratory burst oxidase homolog D (*rbohD*) mutant in Arabidopsis (Zandalinas et al., 2021), no such differences were observed in *C. reinhardtii* between WT and the *rbo1* mutant (Figure 1). Changes in total chlorophyll content of the WT and *rbo1* cells, in response to MFSC, displayed a similar pattern to that shown for growth rate (Figures 1B, S1), as well as to that found for chlorophyll content in Arabidopsis seedlings subjected to MFSC (Zandalinas et al., 2021), supporting the results obtained for *C. reinhardtii* growth under conditions of MFSC (Figures 1A, 1B, S2).

**Figure 1.**
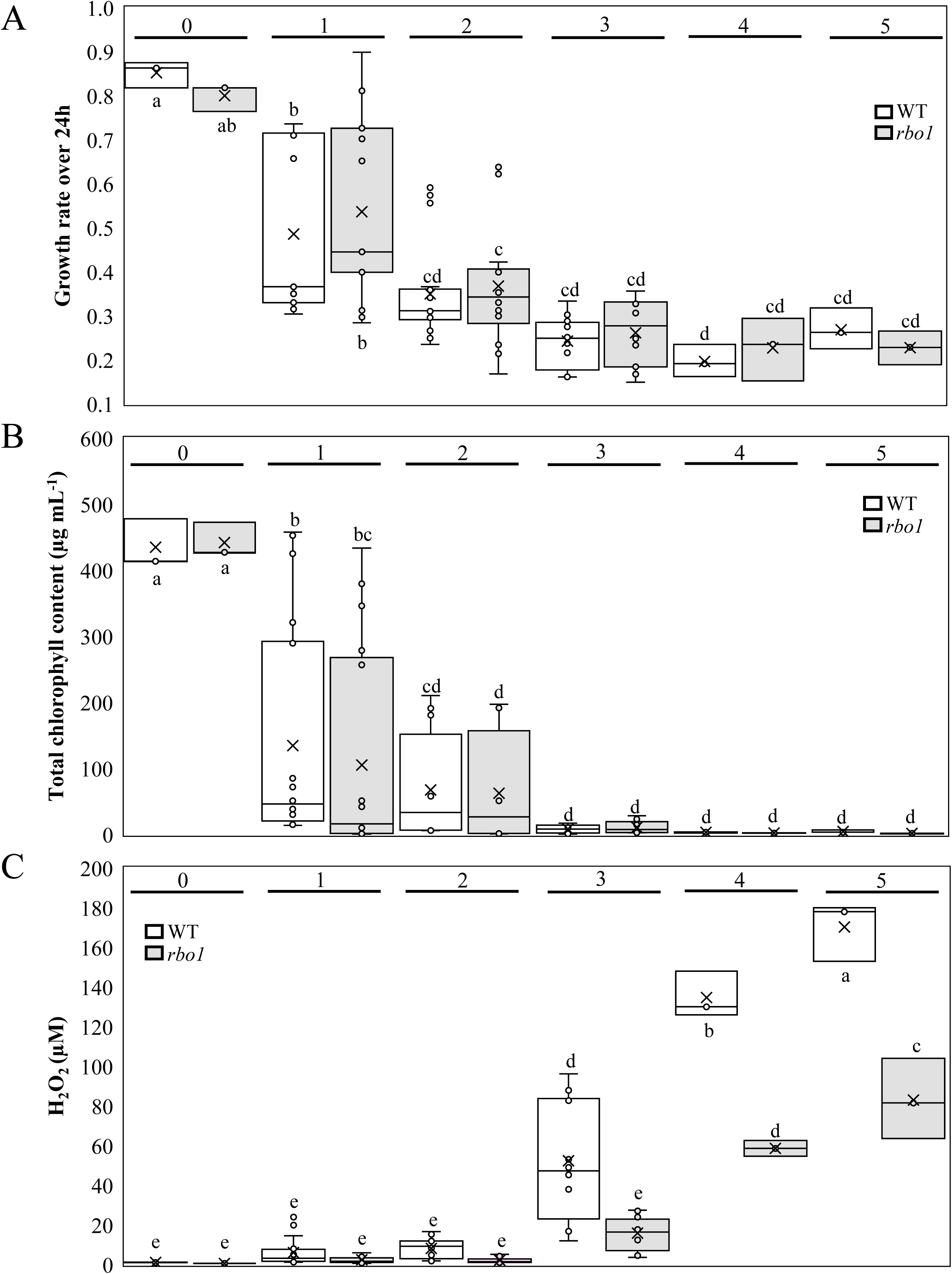
The response of *C. reinhardtii* to a multifactorial stress combination of five different abiotic stresses. **A)** and **B)** The negative impacts of multifactorial stress combination (MFSC) on the growth rate (A) and chlorophyll content (B) of wild type (WT) and the respiratory burst oxidase homolog 1 (*rbo1*) mutant of *C. reinhardtii* grown in liquid cultures. Salinity (S; 120 mM), cadmium (Cd; 300 µM), excess light (EL; 700 μmol m^−2^ s^−1^; a 7-fold increase over control growth conditions), acidic pH (pH; pH 5.5), and the herbicide paraquat (PQ; 50 nM) were used to generate 1-, 2-, 3-, 4-, and 5-stress combinations. **C)** Hydrogen peroxide (H_2_O_2_) levels in the media of WT and *rbo1* cells subjected to the MFSC conditions shown in A and B. All experiments were repeated in 3 biological repeats, each with at least 3 technical repeats. In the graphs, 1 represents an average of all individual stresses (S, Cd, EL, pH, PQ), 2 represents an average of all 2-stress combinations (S+Cd, S+EL, S+PQ, Cd+EL, Cd+PQ, EL+PQ), 3 represents an average of all 3-stress combinations (Cd+EL+S, EL+S+PQ, Cd+EL+PQ, Cd+ S+PQ), 4 represents the 4-stress combination (Cd+EL+S+PQ), and 5 is the 5-stress combination (Cd+EL+S+PQ+pH). Statistical analysis was performed by 2-way ANOVA followed by a Fisher post hoc test (different letters denote statistical significance at p < 0.05).

As WT and the *rbo1* mutant are expected to differ in the levels of ROS they produce during stress (Fichman et al., 2023; Zhou et al., 2024), we measured the H_2_O_2_ concentration in the media of WT and *rbo1* cells subjected to MFSC. As shown in Figure 1C, compared to WT and *rbo1* cells subjected to control, 1-, or 2-stress combinations, the levels of H_2_O_2_ in the media of WT cells increased significantly under the 3-stress combinations, and the levels of H_2_O_2_ increased significantly in WT and *rbo1* under the 4- and 5-stress combinations (Figures 1C, S1). As expected from a *rboh* mutant (Mitter et al., 2022), the levels of H_2_O_2_ found in the media of *rbo1* cells were significantly lower than those in the media of WT cells, subjected to 3-, 4-, and 5-stress combinations (Figure 1C). H_2_O_2_ found in the media of cells could originate from H_2_O_2_ produced inside the cell (*e.g.,* produced in the chloroplast or mitochondria and released into the media through peroxiporins or simple diffusion), or it could be a result of RBOHs found on the plasma membrane of cells (Mittler et al., 2022; Smirnoff and Arnaud, 2019). Further studies are needed to determine the exact source of the H_2_O_2_ found in the media of WT and the *rbo1* mutant during MFSC. The finding that H_2_O_2_ levels in the media of the *rbo1* mutant were reduced (Figure 1C) suggests nonetheless that at least part of the H_2_O_2_ found in the media of *C. reinhardtii* cells during MFSC is generated by RBO1 and the superoxide radicals produced by this enzyme dismutate into H_2_O_2_, spontaneously or enzymatically via superoxide dismutases (SODs; Mittler et al., 2022; Smirnoff and Arnaud, 2019).

### Enhanced cell aggregation in the *rbo1* mutant during MFSC

While subjecting *C. reinhardtii* cells to MFSC for 24 hours, we observed that cells grown at high levels of stress combinations tended to form aggregates (Figure S2). We therefore quantified the number of single cells and cell aggregates in WT and *rbo1 C. reinhardtii* cells in response to MFSC. As shown in Figure 2 and S1, compared to WT cells, *rbo1* cells tended to aggregate even in the absence of stress. In response to 1- and 2- stress combinations, WT aggregated to similar levels as the *rbo1* mutant (significantly higher than WT under control conditions). In contrast, in response to 3- and 4- stress combinations, both WT and *rbo1* had a higher aggregation content compared to control conditions and 1- and 2-stress combinations. In response to 5-stress combinations, that included acidic pH, a known inducer of aggregation in *C. reinhardtii*, both WT and *rbo1* cells displayed the highest level of aggregations, with *rbo1* having significantly more aggregation compared to WT (Figures 2, S1). As aggregation in *C. reinhardtii* is typically associated with stress (Ratcliff et al., 2013; de Carpentier et al., 2019), our findings could suggest that under control conditions, or in response to the 5-stress combination conditions, *rbo1* cells experience more stress compared to WT. While this result is interesting, and perhaps expected, the findings that WT and *rbo1* did not differ in their growth rate and chlorophyll content during MFSC (Figure 1) could suggest that aggregation is a result of differences in inter- or intra-cellular H_2_O_2_ levels or media pH. We therefore tested the effect of extracellular H_2_O_2_ application on aggregation of WT and *rbo1* cells in the absence of stress. As shown in Figure 3A-3B, while addition of low concentrations of H_2_O_2_ (*i.e.,* 0.5 or 1.5 µM) to the media prompted similar aggregation in WT and *rbo1*, compared to WT, *rbo1* cells produced more aggregates at higher H_2_O_2_ levels of 3 or 5 µM.

**Figure 2.**
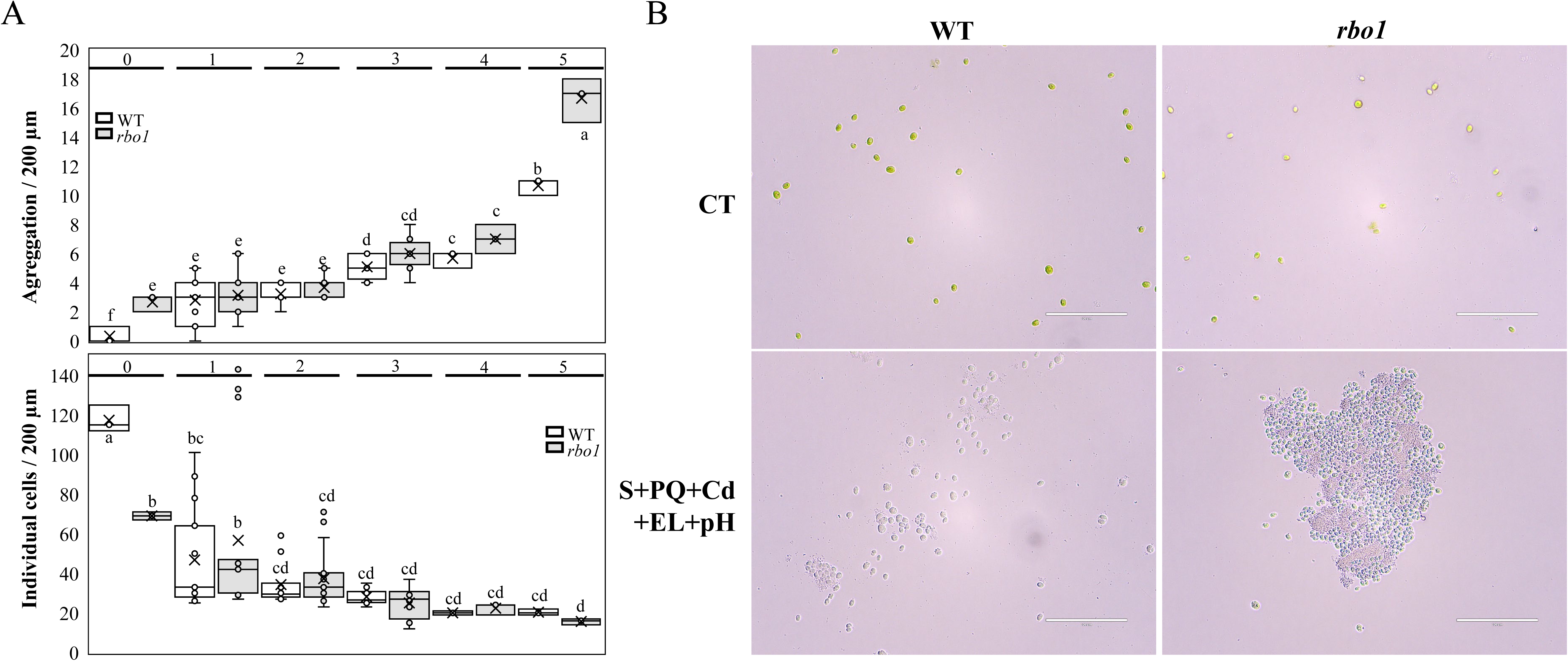
Enhanced aggregation in wild type and the *rbo1* mutant in response to a multifactorial stress combination of five different abiotic stresses. **A)** Enhanced number of aggregates (top) and decreased number of individual cells (bottom) in wild type (WT) and the respiratory burst oxidase homolog 1 (*rbo1*) mutant of *C. reinhardtii* subjected to multifactorial stress combination (MFSC; as described in Figure 1). **B)** Representative images of WT and *rbo1* cells grown under control (CT) conditions or subjected to a MFSC of salinity (S), cadmium (Cd), excess light (EL), acidic pH (pH), and (PQ) combined (5-stress MFSC). All experiments were repeated in 3 biological repeats, each with at least 3 technical repeats. Statistical analysis was performed by 1-way ANOVA followed by a Fisher post hoc test (different letters denote statistical significance at p < 0.05). Bars in images indicate 100 µm.

**Figure 3.**
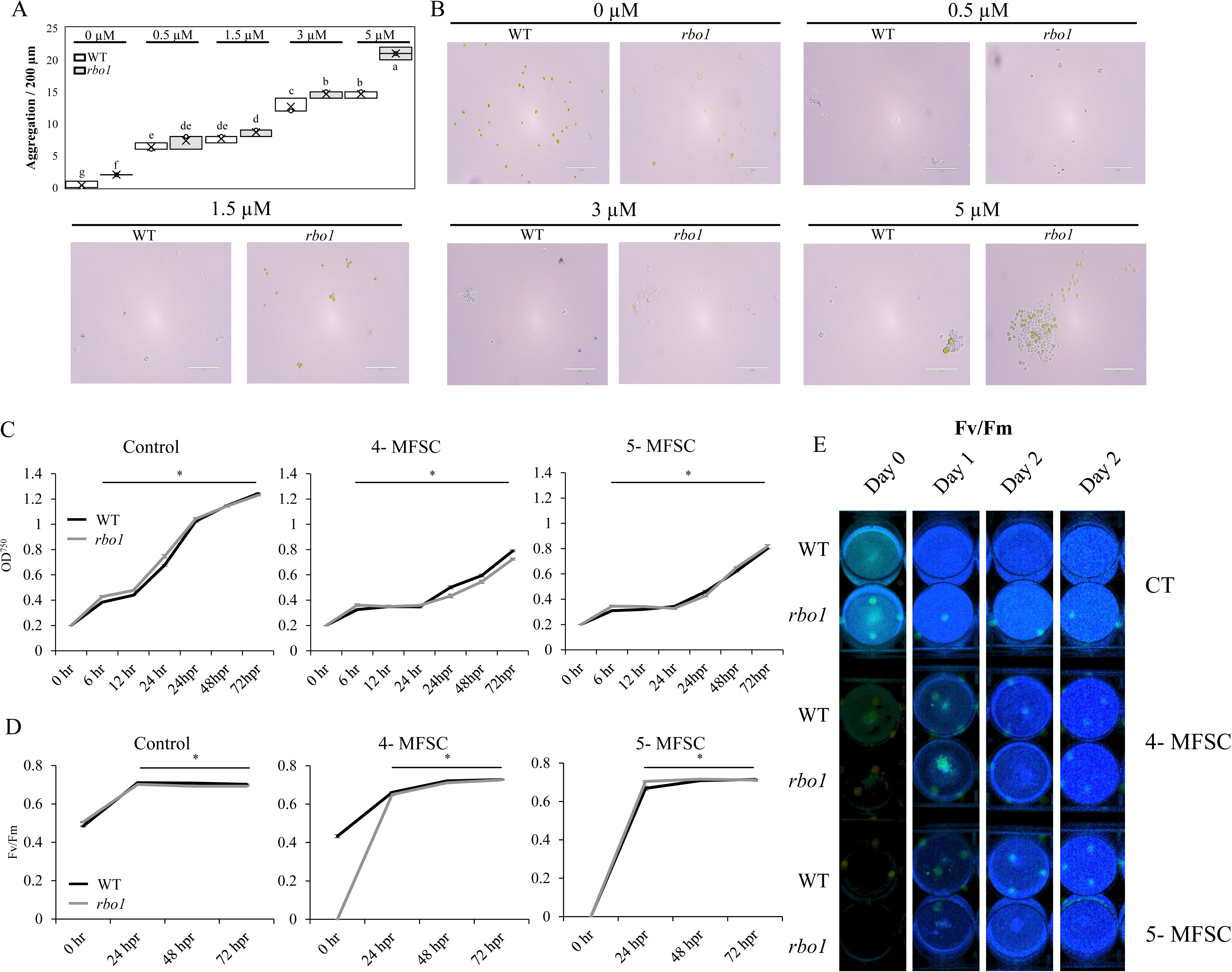
Enhanced aggregation in *C. reinhardtii* wild type and *rbo1* cells in response to exogenous H_2_O_2_ application, and recovery from MFSC. **A)** Enhanced number of aggregates in wild type (WT) and the respiratory burst oxidase homolog 1 (*rbo1*) mutant of *C. reinhardtii* treated with different concentrations of H_2_O_2_. **B)** Representative images of WT and *rbo1* cells grown under control (CT) conditions or subjected different concentrations of H_2_O_2_. **C)** Recovery from the 24 hour 4- and 5-MFSCs (Cd+EL+S+PQ, Cd+EL+S+PQ+pH, respectively) measure by culture OD_750_. **D)** Same as in C, but measured by the ratio between variable and maximum fluorescence (Fv/Fm). **E)** Representative images of *C. reinhardtii* cultures imaged for Fv/Fm. All experiments were repeated in 3 biological repeats, each with at least 3 technical repeats. Statistical analysis was performed by 2-way ANOVA followed by a Fisher post hoc test (different letters denote statistical significance at p < 0.05), or by two-tailed Student’s t-test (asterisks denote statistical significance at p < 0.05 compared to time 0 hour). Bars in images indicate 100 µm.

Taken together, the results shown in Figures 2 and 3A-3B suggest that although the *rbo1* mutant accumulated less H_2_O_2_ in its growth media, it was more sensitive to H_2_O_2_ and formed cell aggregates in response to lower levels of H_2_O_2_. This result could suggest that the *rbo1* mutant is more sensitive to H_2_O_2_ due to higher sensing of H_2_O_2_, or lower ability to scavenge H_2_O_2_ (a potential result of acclimating to a lower production rate of H_2_O_2_; Mittler et al., 2022).

To test whether *C. reinhardtii* cells subjected to the MFSC treatments can recover, we subjected WT and *rbo1* cells to the 4- or 5- stress MFSC treatments (S+Cd+EL+PQ or S+PQ+Cd+EL+pH, respectively) for 24 hours, and allowed them to recover. As shown in Figure 3C-3E, *C. reinhardtii* cells subjected to these MFSC treatments were able to recover and grow, as well as recover some of their photosynthetic parameters. These findings reveal that although *C. reinhardtii* cells stopped growing and aggregated during MFSC, at least under the conditions we used, they did not die and were able to recover.

To further dissect the responses of WT and the *rbo1* mutant to MFSC and study the expression of stress-response and ROS-metabolizing enzymes in these two genotypes subjected to MFSC, we conducted a whole-cell proteomics analysis of WT and *rbo1* subjected to MFSC.

### Proteomics analysis of *C. reinhardtii* response to MFSC

Whole-cell proteomics analysis was conducted using WT and *rbo1 C. reinhardtii* cells subjected to the same MFSC conditions shown in Figures 1 and 2 (Figures 4, 5, S3-S8; Tables S1-S46). While the response of *C. reinhardtii* to each of the individual stresses applied resulted in the identification of pathways and genes that matched each stress (*e.g.,* Fe-S proteins in Cd, oxidoreductases in PQ, ion channels in S, photosynthesis in EL, and ATPase-coupled transporters in pH; Figure S3, Tables S45 and S46), we found no overlapping proteomic responses shared by all 5 individual stresses (Figure 4A). When different stresses were combined at the 2-stress level, we found shared responses between PQ+EL and Cd+PQ, but fewer shared responses between other stresses (Figure 4B). At the 3-, 4- and 5-stress combination level, similarities in responses were mainly observed between different combinations that included S, Cd, EL, PQ, and pH, but this similarity was also low (Figure 4C). Pathways identified in WT cells subjected to the 4- and 5- stress combinations included amino acid and pigment biosynthesis and degradation, motility, cytoskeleton organization, and response to chemicals (Figure 4D). Some of these corresponded with the increase in cell aggregation under MFSC (*e.g.,* enhanced motility, membrane structures, *etc.*). Proteomics analysis of the *rbo1* mutant revealed that, even in this mutant, there were no overlapping proteomic responses shared by all 5 individual stresses (Figure S4). The majority of proteins found to differ between WT and the *rbo1* mutant in responses to the different stresses (Figures S4, S5) were involved in cell motility and metabolic processes, however, the major stress-response groups of WT and *rbo1* were overall similarly expressed (Figures 4, 5A, S3-S6). In response to the 4- and 5-stress combinations, and in agreement with the cell aggregation results (Figures 2, 3), the abundance of proteins associated with motility, membrane and extracellular organization, and cytoskeleton/cilium functions was also high in the *rbo1* mutant (Figures 4, 5A, S3-S6).

**Figure 4.**
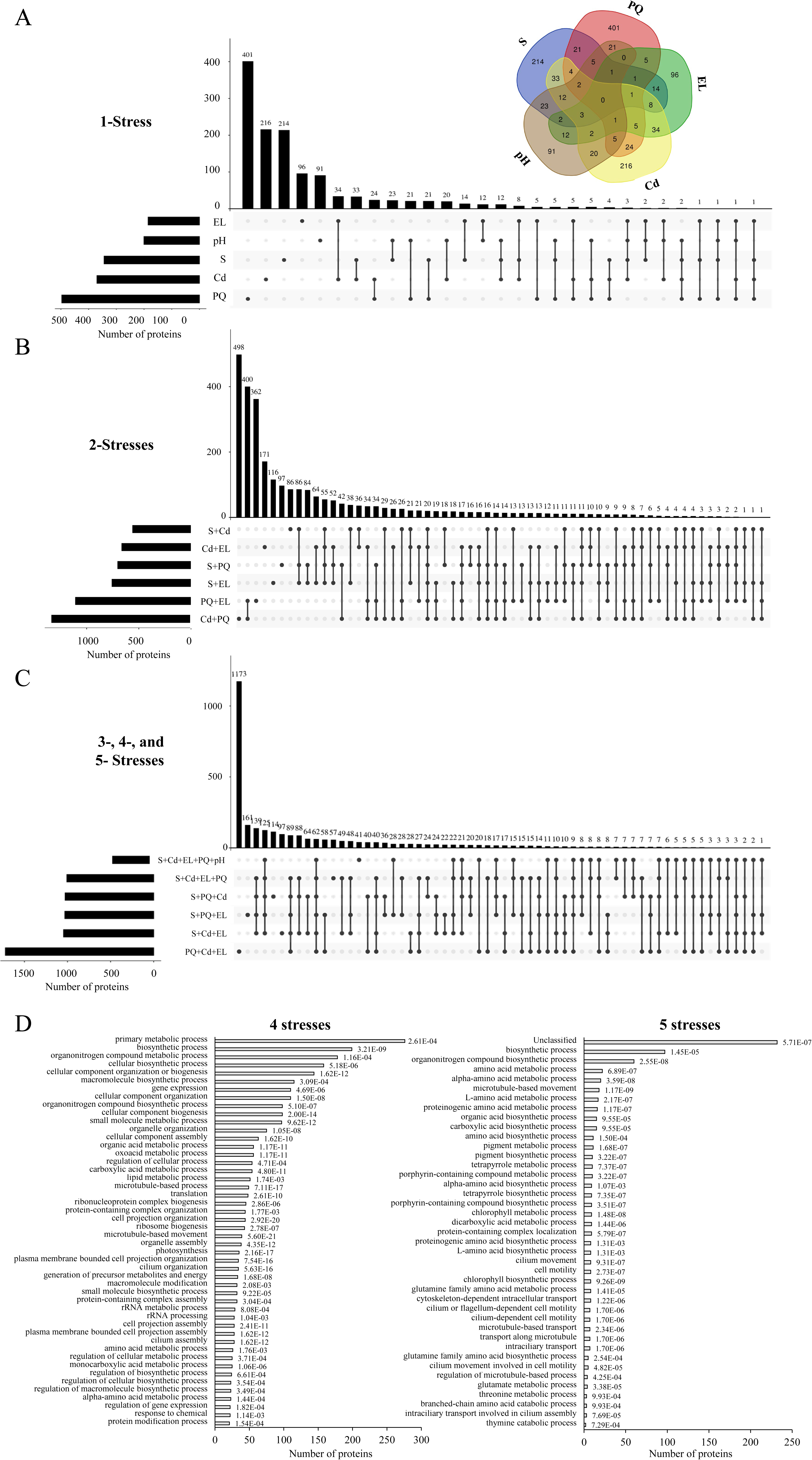
Proteomics analysis of wild type *C. reinhardtii* cells subjected to a multifactorial stress combination of five different abiotic stresses. **A)** Venn (top) and upset (bottom) plots showing the overlap between different proteins with a significant change in abundance in response to treatment with salinity (S), cadmium (Cd), excess light (EL), acidic pH (pH), or paraquat (PQ). **B)** and **C)** Upset plots showing the overlap between different proteins with a significant change in abundance in response to all 2-stress combination treatments (B), or all 3-, 4-, and 5-stress combination treatments (C). **D)** Gene ontology (GO) annotation of proteins with a significant change in abundance in response to 4- (left) or 5- (right) stress combinations.

As RBO1 is associated with ROS production in cells (Fichman et al., 2023; Zhou et al., 2024), we compared the abundance of ROS-metabolizing proteins between WT and *rbo1* cells subjected to MFSC (Figures 5B, S7). This analysis revealed that while the ROS scavenging enzyme glutathione peroxidase 5 (GPX5; Ma et al., 2020; Youssef et al., 2023) increased in abundance in WT during the 4- and 5-stress combinations, the level of this protein did not change in the *rbo1* mutant under the same conditions (Figure 5B). In contrast, the abundance of mitochondrial cytochrome c-peroxidase increased in the *rbo1* mutant, but not WT under the 5-stress combination (Figure 5B). Interestingly, many of the other major H_2_O_2_ scavenging enzymes, such as catalase, peroxiredoxin, and ascorbate peroxidase, did not significantly increase in their abundance in response to the 4- and 5-stress combinations, while manganese superoxide dismutase (MSD2 and/or MSD3) significantly increased in abundance in both WT and the *rbo1* mutant under the same conditions (Figure 5B). Under non stress conditions, *rbo1* cells accumulated a thioredoxin-dependent peroxiredoxin isoform that was not accumulated in WT (Figure S7B). Taken together, the findings presented in Figures 1C, 2, 3, 5B and S7 suggest that under a combination of multiple stresses (especially the 4- and 5-stress combinations) *rbo1* cells produce less H_2_O_2_ due to the lack of RBO1, but are more sensitive to H_2_O_2_, and aggregate more, possibly due to the suppressed expression of GPX5. Further studies are required to determine the exact subcellular distribution of different ROS in WT and *rbo1* cells under the different stress conditions, as well as the function of all main ROS producing and scavenging enzymes of *C. reinhardtii* WT and *rbo1* cells under conditions of MFSC.

**Figure 5.**
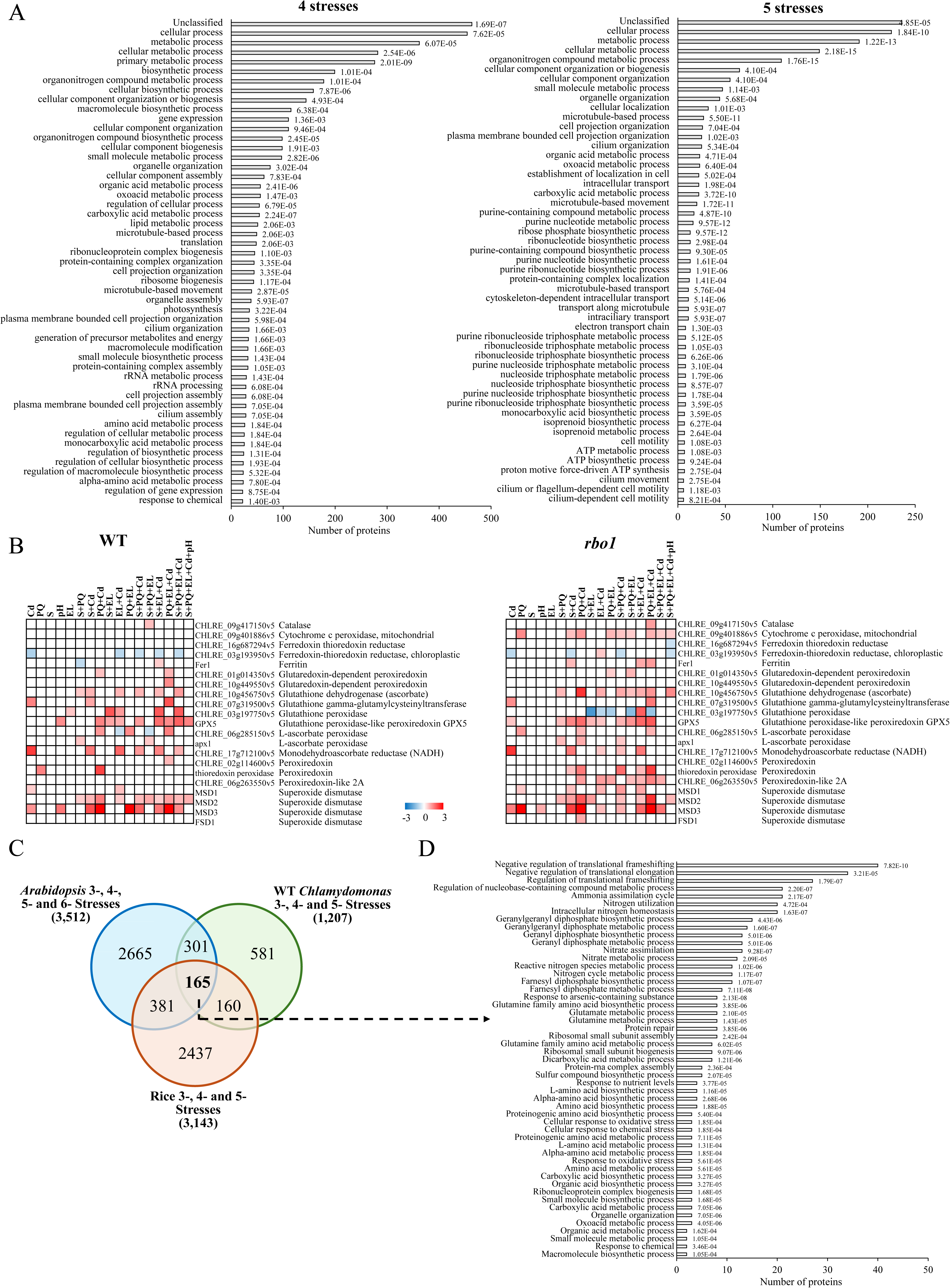
Proteomics analysis of *rbo1 C. reinhardtii* cells subjected to multifactorial stress combination (MFSC) and a comparison between the responses of *Arabidopsis thaliana* and *C. reinhardtii* to MFSC. **A)** Gene ontology (GO) annotation of proteins with a significant change in abundance in response to 4- (left) or 5- (right) stress combinations in the respiratory burst oxidase homolog 1 (*rbo1*) mutant. **B)** Partial heatmaps of reactive oxygen species (ROS) metabolizing enzymes abundance in WT (left) and *rbo1* (right) cells subjected to a MFSC of salinity (S), cadmium (Cd), excess light (EL), acidic pH (pH), and/or paraquat (PQ). Complete heatmaps are shown in Figure S7. **C)** Venn diagram showing the overlap between *Arabidopsis* homologs of *C. reinhardtii* wild type proteins significantly altered in their abundance in response to a MFSC of 3-, 4-, and 5-stress combinations, *Arabidopsis* homologs of rice proteins significantly altered in their abundance in response to 3-, 4-, and 5-stress MFSC, and *Arabidopsis* transcripts significantly altered in their expression in response to a MFSC of 3-, 4-, 5-, and 6-stress combinations. **D)** GO analysis of the 165 Arabidopsis homologs common to the response of *C. reinhardtii*, *A, thaliana*, and rice to MFSC (from C).

To compare between the responses of *C. reinhardtii*, *A. thaliana* (Zandalinas et al., 2021), and rice (Sinha et al., 2024) to MFSC, we compiled a list of all proteins altered in their abundance in *C. reinhardtii* WT in response to the 3-, 4- and 5-MFSCs (Table S35). We then used the Ensembl BioMart Homologues and OMA Browser genome pair orthology tools (https://www.ensembl.org/biomart; Altenhoff et al., 2021) to identify their *A. thaliana* homologs (1,207; Tables S36). A similar analysis was done to determine the *A. thaliana* homologs of rice proteins altered in their response to MFSC (3,143; Sinha et al., 2024; Table S37). We then determined the overlap between the lists of *C. reinhardtii* proteins (1,207; Table S38), Arabidopsis transcripts (3,512; Zandalinas et al., 2021; Peláez-Vico et al., 2024; Table S40), and rice proteins (3,143 proteins in rice; Sinha et al., 2024; Table S39), altered in response to MFSC. As shown in Figure 5C, 165 homologs were found to be common to the responses of *A. thaliana,* rice, and *C. reinhardtii* to MFSC (Table S41). A GO analysis of these 165 homologs identified transcripts involved in nitrogen, geranylgeranyl diphosphate, glutamine, and oxoacid metabolism, as well as nutrient stress and reactive nitrogen and oxygen species, as enriched in this conserved group (Figures 5D, S8). Most of these transcripts were not previously identified as involved in MFSC and their function should be addressed in future studies.

### Summary

Previous studies of MFSC focused on flowering plants, which are multicellular photosynthetic organisms. In contrast, the work presented here focused on a unicellular photosynthetic organism (*C. reinhardtii*). While several similarities were identified between the responses of multicellular and unicellular organisms to MFSC (*e.g.,* the impact of MFSC on growth, chlorophyll content, ROS accumulation, and the increased abundance of Cu, Fe, and Fe-S binding proteins; Figures 1, 4, 5; Table S38), our work revealed a novel response to MFSC, *i.e.,* enhanced cell aggregation (Figure 2). We further show that aggregation in response to MFSC is elevated in the *rbo1* mutant, which is impaired in ROS metabolism, and that it can be induced by external application of H_2_O_2_ (Figures 2, 3). Aggregation is considered an ancient form of multicellularity and is induced in many unicellular organisms by stress (Márquez-Zacarías et al., 2021; Tong et al., 2022). Our findings that aggregation is triggered in *C. reinhardtii* by MFSC and is associated with ROS metabolism (Figures 2-5), could shed new light on the response of unicellular organisms to MFSC, and suggest that one of the main drivers leading to multicellularity during evolution was early conditions of MFSC under an oxygenated environment. As aggregation on solid surfaces can reduce growth, respiration and/or photosynthesis (due *e.g.,* to diffusion barriers; Márquez-Zacarías et al., 2021; Tong et al., 2022), enhanced aggregation of unicellular organisms in different ecosystems subjected to MFSC could also explain part of the negative impacts MFSC has on the services provided by these complex biosystems (Rillig et al., 2019; Rillig et al., 2021; Speißer et al., 2022). Further studies are needed to determine the negative effects of different types of MFSC (including for example heat stress) on different unicellular organisms across ecosystems worldwide, the role of aggregation in these negative impacts, and the relationships between these processes and anthropogenic activities.

## Materials and Methods

### Organisms, stress treatments, growth rate, and chlorophyll content

*C. reinhardtii* WT and *rbo1* (lmj.ry0402.234817 from the Chlamydomonas Resource Center, University of Minnesota, St. Paul, MN, USA, and UTEX Culture Collection of Algae; Fichman et al., 2023; Zhou et al., 2024) were grown in liquid TAP media, pH 7, 21^°^C, 8 h/16 h light/dark, 100 μmol m^−2^ s^−1^, and 150 rpm. Stress conditions included excess light (700 μmol m^−2^ s^−1^; EL), paraquat (50 nM; PQ), salinity (120 mM NaCl; S), heavy metal (Cd, 300 μM CdCl_2_), and acidity (pH = 5.5). Cd, EL, S, and PQ were applied in all possible combinations, and pH was added as single stress, as well as in combination with Cd+EL+S+PQ to generate five-stress combination (Pascual et al., 2023; Peláez-Vico et al., 2023; Sinha et al., 2024) in 6-wells plate using a media volume of 4 mL. *C. reinhardtii* WT and *rbo1* were inoculated at an initial OD_750_ of 0.2 and subjected to the different stresses for 1 day. OD_750_ was assessed at 6, 12 and 24 h using a Cytation 5 imaging plate reader (Agilent, USA) as described in Bernd and Cook (2002). Measurements were taken after 30 seconds of shaking and the growth rate was calculated based on the change in OD over time. pH was measured before and after applying stressors to the media. Chlorophyll content was quantified as described in Zhang et al., (2022). Three different replicates for each genotype and condition were measured.

### Microscopy

Cells from each condition and genotype were mounted on slides and imaged using an EVO XL Cell Imaging System Amex3300 Inverted Digital Microscope (Thermo Scientific, Massachusetts, USA). Images were taken at 20X and 40X and the number of aggregated *Chlamydomonas* cells (defined as three or more cells in close proximity) and individual cells (cells exhibiting no palmelloid formation) were counted. A minimum of 20 images were taken for each condition. To determine the role of H_2_O_2_ in aggregation (Figure 3), H_2_O_2_ (0, 0.5, 1.5, 3 and 5 µM) was applied to the TAP media and cells were grown as described above for 24 h before images were taken.

### H_2_O_2_ concentration

H_2_O_2_ in the growth media was quantified using Amplex-Red (10-Acetyl-3,7-dihydroxyphenoxazine; Thermo Fisher Scientific, Waltham, Massachusetts, USA). 2 mL of culture was centrifuged for 10 minutes at 12,000 g, 4°C, and the supernatant was recovered and buffered with 1 M phosphate buffer pH 7.4. H_2_O_2_ quantification was performed according to the MyQubit-AmplexRed Peroxide Assay manual (Thermo Fisher Scientific), using an H_2_O_2_ calibration curve (Thermo Fisher Scientific).

### Recovery experiments and photosynthetic measurements

To determine whether WT and *rbo1* cells can recover from the 24 hour 4- or 5- MFSC treatments (S+PQ+Cd+EL, or S+PQ+Cd+EL+pH, respectively), stress and control treated cells were centrifuged for 10 min at 3,000 g at the end of the stress treatment, resuspended in TAP media, pH 7, and kept at 21^°^C, 8 h/16 h light/dark, 100 μmol m^−2^ s^−1^, and 150 rpm for 72 hours. OD_750_ was assessed at 0, 6, 12, 24, 48 and 72 h post stress using a Cytation 5 imaging plate reader (Agilent, USA). In addition, the ratio of variable fluorescence to maximum fluorescence (Fv/Fm; Hemker et al., 2024) was measured using a Hexagon imaging PAM (HEXAGON-IMAGING-PAM, Walz, Effeltrich, Germany) apparatus at 0 (following the end of the 24 hour stress treatments), and at 24, 48, and 72 hours of recovery, according to manufacturer instructions.

### Proteomics Analysis

Cultures (4 mL) of cells grown and stressed as described above for 24 h were centrifuged for 10 minutes at 12,000 g, 4°C. Pellets were resuspended in 1X Laemmli buffer, boiled, and centrifuged at 16,000 g for 10 min. The supernatant was precipitated with cold acetone. The resulting protein pellets were resuspended in a solution containing 6 M urea, 2 M thiourea, and 100 mM ammonium bicarbonate. Protein concentration was quantified using the Pierce 660 nm Protein Assay. 20 µg of protein from each sample were reduced, alkylated, and digested by trypsin as described in Sinha et al., (2024). 500 ng of digested peptides were subjected to proteomics analysis as described in Sinha et al., (2024) using an Evosep One LC system interfaced with a Bruker timsTOF PRO2 mass spectrometer (Bruker) using the *C. reinhardtii* UniProt database (accession UP000006906), which comprises 18,829 protein entries. A false discovery rate of 1% was set for peptide spectrum match, peptide, and protein identification. Protein quantification was performed using MaxLFQ based on the area of MS2, with cross-run normalization enabled and no imputation applied. Differential abundance testing was conducted using unpaired t-tests, and significant candidates were identified based on a significance threshold of p-value < 0.01, q-value < 0.05, and an absolute average log_2_ (fold change) > 0.58. For all proteins studied, the reference was the expression value obtained for the CT conditions.

### Statistical Analysis

All experiments were repeated in 3 biological repeats, each with at least 3 technical repeats. Box plots are presented with mean as X ± SE; median is the line in the box and box borders are 25^th^ and 75^th^ percentiles; whiskers are the 1.5 interquartile range. Statistical analysis was performed with Statgraphics Plus v.5.1. software (Statistical Graphics Corp., Herndon, VA, United States) by two-way analysis of variance (ANOVA) followed by Fisher’s post hoc test (different letters denote statistical significance at p < 0.05), or by two-tailed Student’s t-test (asterisks denote statistical significance at p < 0.05 compared to time 0h).

## ACKNOWLEDGMENTS

This work was supported by funding from the National Science Foundation (IOS- 2414183; IOS-2110017, IOS-2343815), the Interdisciplinary Plant Group, University of Missouri, Columbia, MO, project PID2021-128198OA-I00 funded by Ministerio de Ciencia e Innovación/Agencia Estatal de Investigación MCIN/AEI/10.13039/501100011033 and FEDER Una manera de hacer Europa, and Ramón y Cajal program (RYC2020-029967-I).LSP was supported by the contract PRE2022-101650 funded by MCIN/AEI/10.13039/501100011033 and FSE^+^.

## AUTHOR CONTRIBUTIONS

L.S.P, R.S, D.M, T.T.N., Z.L, B.P.M. T.J., and L.R. performed experiments and analyzed the data. L.S.P., R.M., F.B.F, A.G.C., and S.I.Z. designed experiments, analyzed the data, and/or wrote the manuscript. R.M., A.G.C., and S.I.Z. provided financial support.

### DATA AVAILABILITY

The data that supports the findings of this study are available in the text, figure, and supplemental material of this article. Proteomics data was deposited in PRIDE (https://www.ebi.ac.uk/pride/), under the following accession number: PXD058190.

## Supplementary Figures

**Figure S1.**
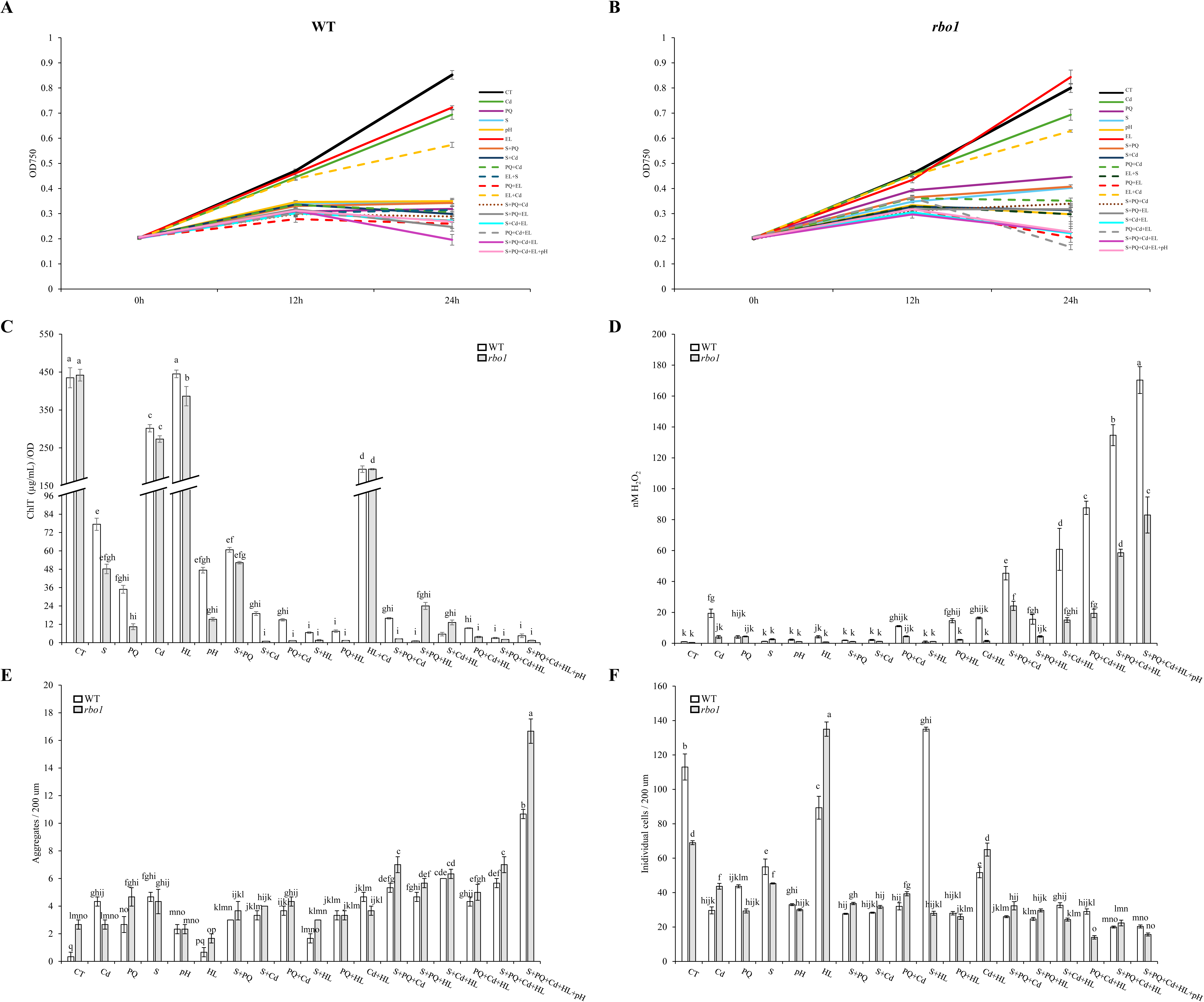
The impacts of multifactorial stress combination (MFSC) on the growth rate **(A, B)**, chlorophyl **(C)** and media H_2_O_2_ **(D)** content, and aggregation **(E, F)** of *C. reinhardtii* wild type (WT) and the respiratory burst oxidase homolog 1 (*rbo1*) mutant cells. Cells were grown in liquid TAP media cultures and subjected to: Salinity (120 mM; S), cadmium (300 µM; Cd), excess light (700 μmol m^−2^ s^−1^; a 7-fold increase over control growth conditions; EL/HL), acidic pH (pH 5.5; pH), and paraquat (50 nM; PQ) in 1-, 2-, 3-, 4-, and 5-stress combinations. All experiments were repeated in 3 biological repeats, each with at least 3 technical repeats. In support of Figures 1 and 2.

**Figure S2.**
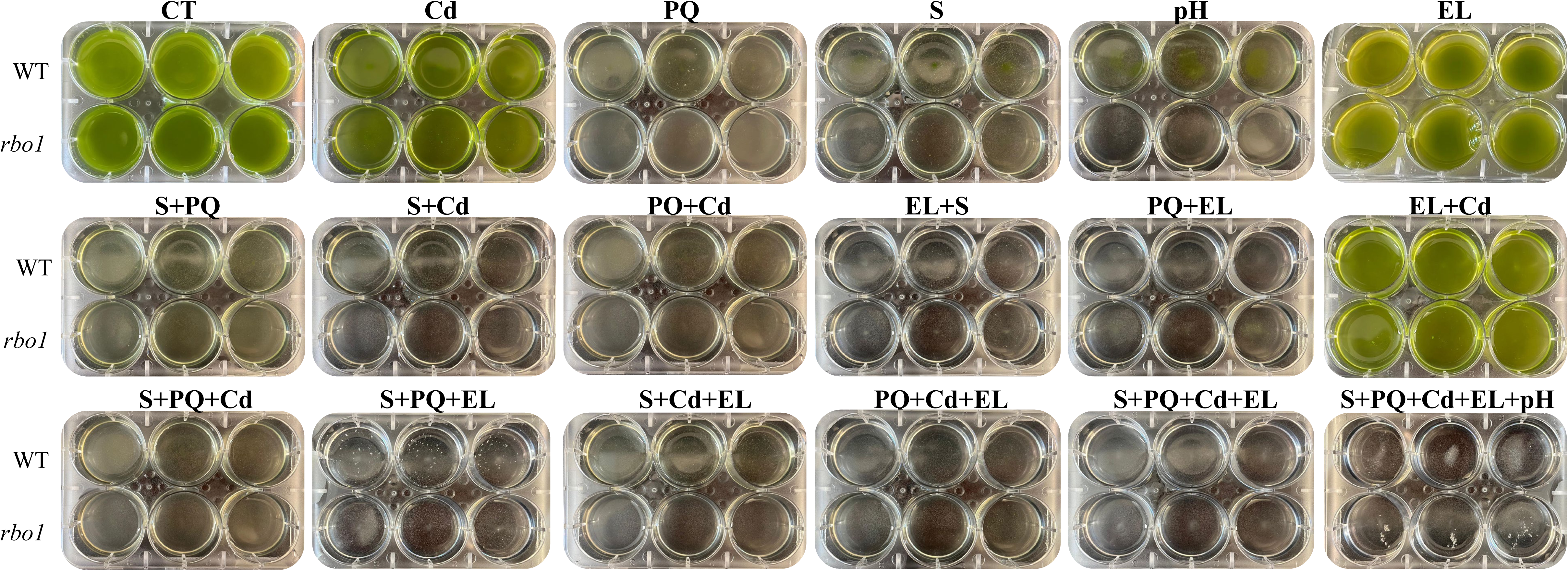
Representative images of *C. reinhardtii* wild type (WT) and respiratory burst oxidase homolog 1 (*rbo1*) cells grown in liquid cultures and subjected to: Salinity (120 mM; S), cadmium (300 µM; Cd), excess light (700 μmol m^−2^ s^−1^; a 7-fold increase over control growth conditions; EL), acidic pH (pH 5.5; pH), and paraquat (50 nM; PQ) in 1-, 2-, 3-, 4-, and 5-stress combinations. All experiments were repeated in 3 biological repeats, each with at least 3 technical repeats. In support of Figures 1 and 2.

**Figure S3.**
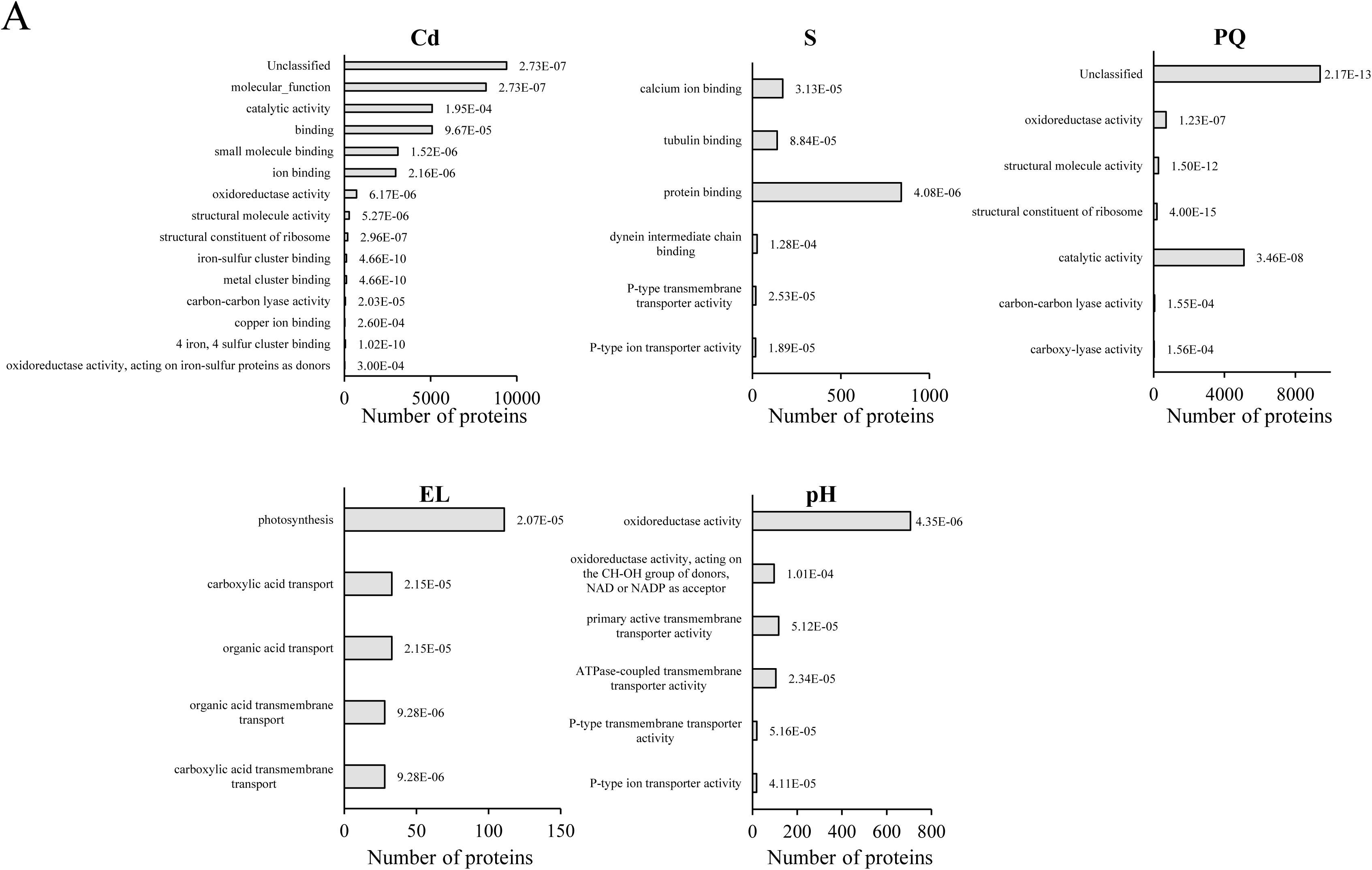
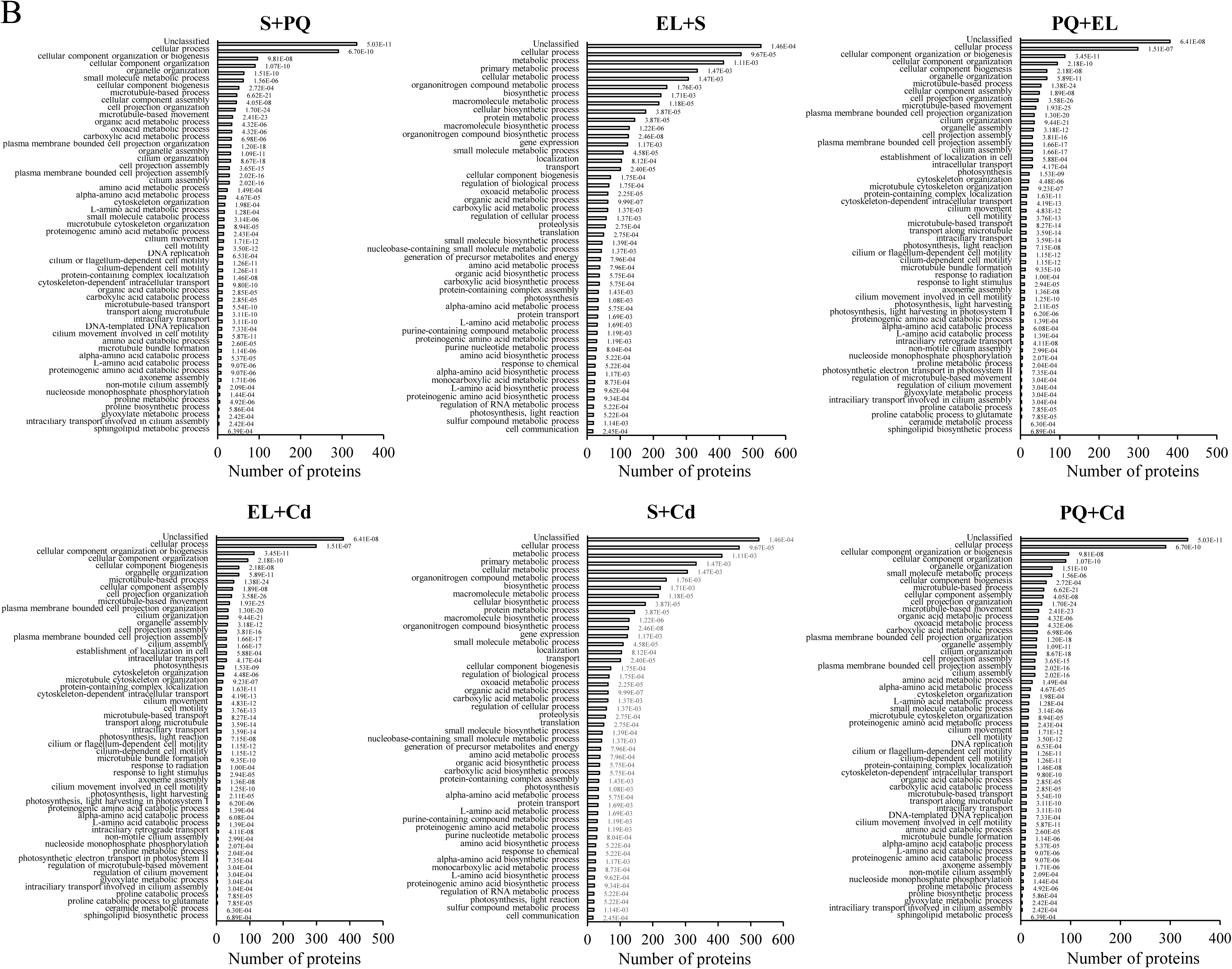
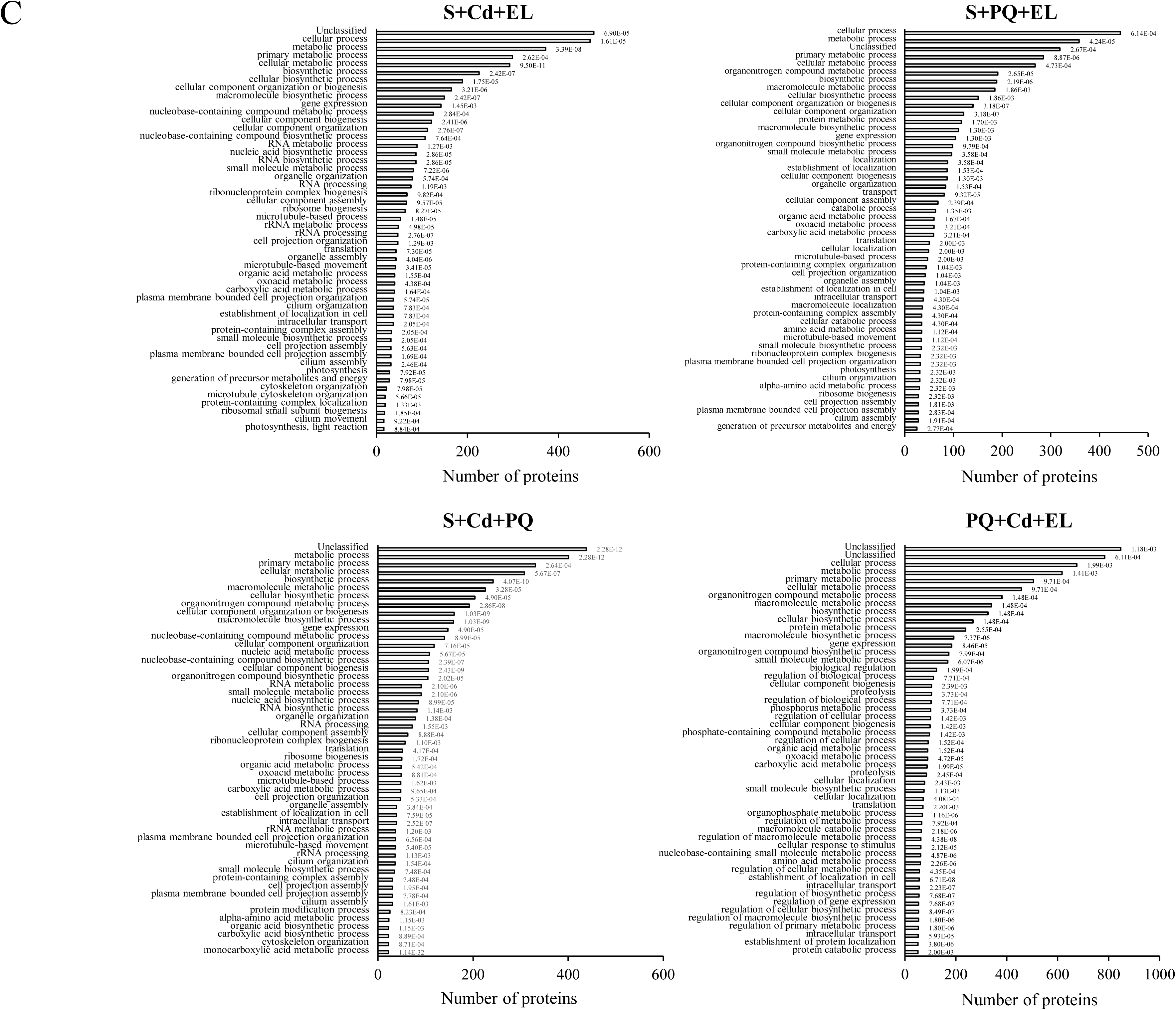
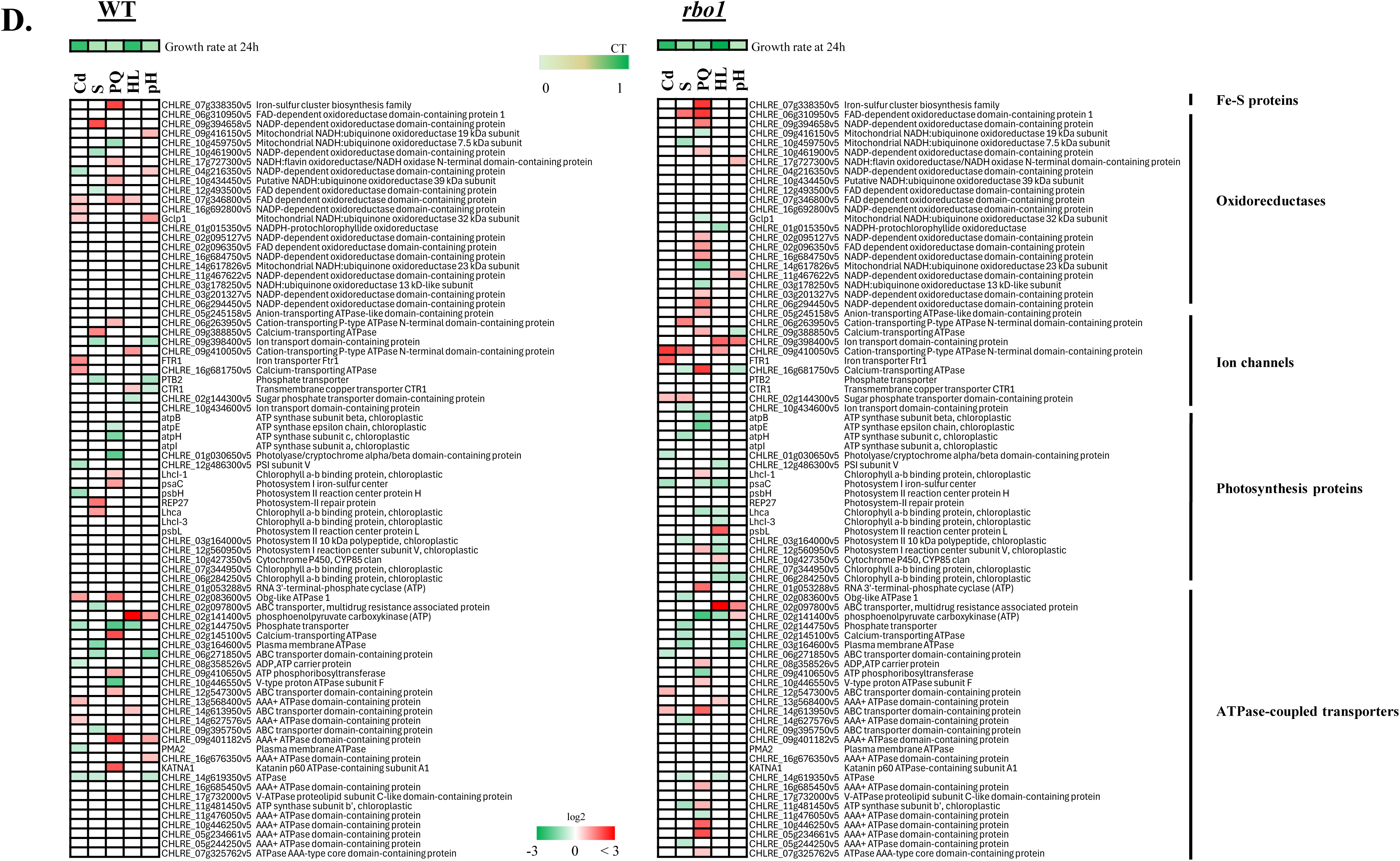
Gene ontology (GO) annotation of wild type (WT) proteins with a significant change in abundance in response to multifactorial stress combination. **A)** GO annotation of proteins with a significant change in abundance in response to treatment with salinity (S), cadmium (Cd), excess light (EL), acidic pH (pH), or paraquat (PQ). **B)** Same as A, but for all 2-stress combinations. **C)** Same as A, but for all 3-, 4-, and 5-stress combinations. **D**) Heatmaps of Fe-S proteins, oxidorrecductases, ion channels, photosynthesis proteins and ATPase-cpupled transporters in WT (left) and *rbo1* (right) *C. reinhardtii* cells subjected to a MFSC of salinity (S), cadmium (Cd), excess light (EL), acidic pH (pH), and/or paraquat (PQ). In support of Figures 4 and 5.

**Figure S4.**
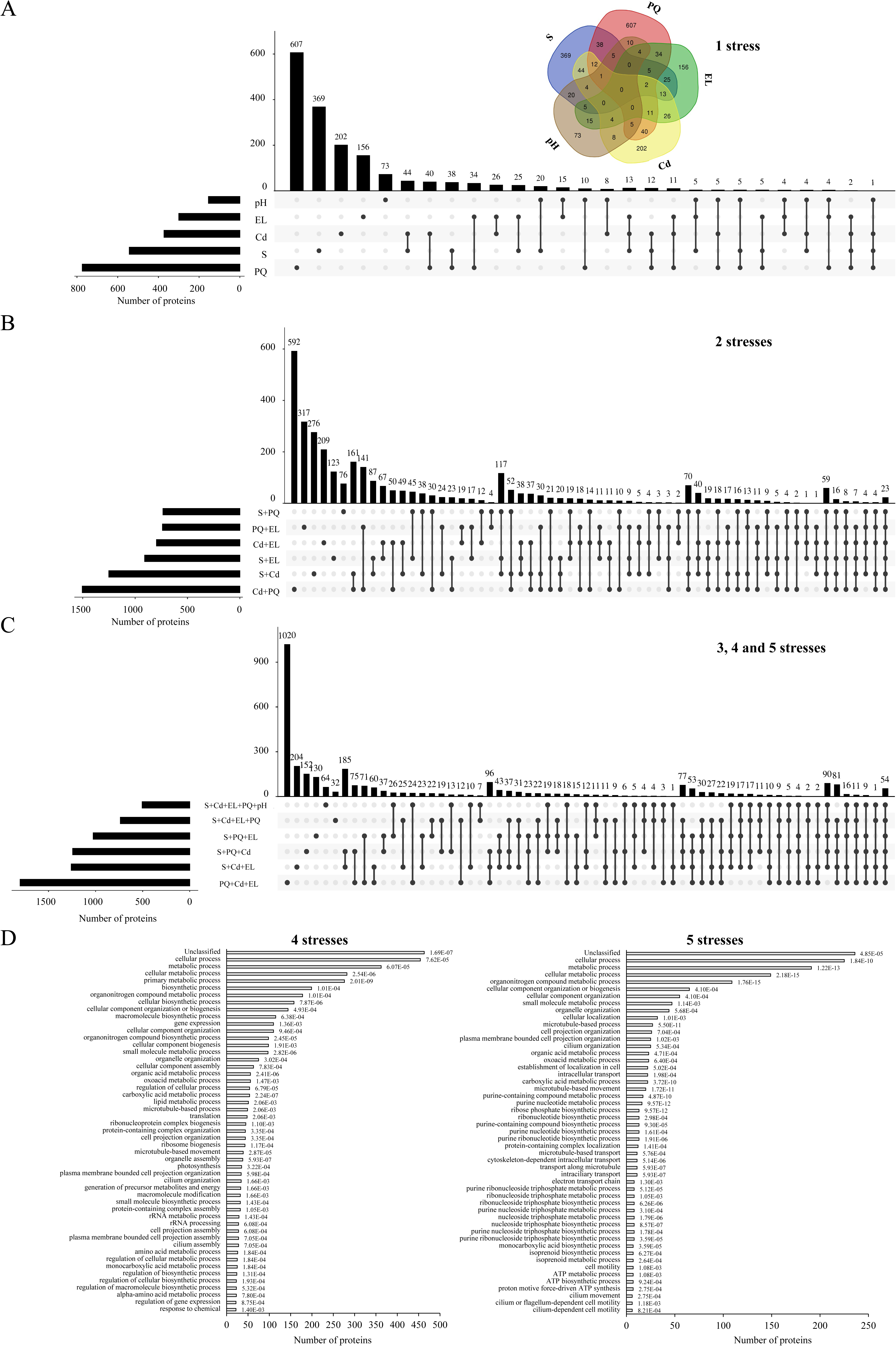
Proteomics analysis of respiratory burst oxidase homolog 1 (*rbo1*) *C. reinhardtii* cells subjected to a multifactorial stress combination of five different abiotic stresses. **A)** Venn (top) and upset (bottom) plots showing the overlap between different proteins with a significant change in abundance in response to treatment with salinity (S), cadmium (Cd), excess light (EL), acidic pH (pH), or paraquat (PQ). **B)** and **C)** Upset plots showing the overlap between different proteins with a significant change in abundance in response to all 2-stress combination treatments (B), or all 3-, 4-, and 5-stress combination treatments (C). **D)** Gene ontology (GO) annotation of proteins with a significant change in abundance in response to 4- (left) or 5- (right) stress combinations. In support of Figures 4 and 5.

**Figure S5.**
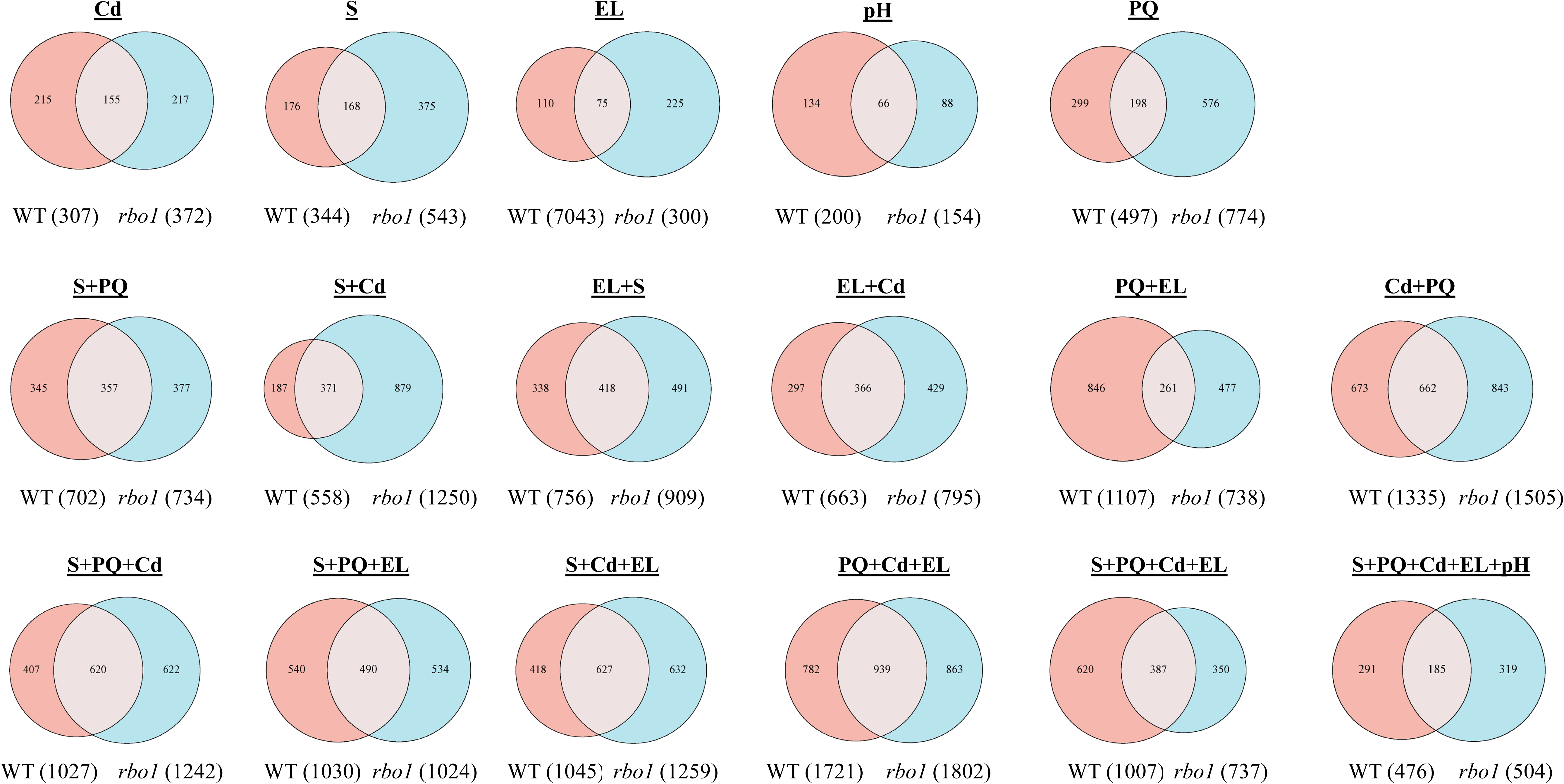
Venn diagrams showing the overlap between proteins with a significant change in abundance in *C. reinhardtii* wild type (WT) and respiratory burst oxidase homolog 1 (*rbo1*) cells subjected to: Salinity (120 Mm; S), cadmium (300 µM; Cd), excess light (700 μmol m^−2^ s^−1^; a 7-fold increase over control growth conditions; EL), acidic pH (pH 5.5; pH), and paraquat (50 nM; PQ) in 1-, 2-, 3-, 4-, and 5-stress combinations. In support of Figures 4 and 5.

**Figure S6.**
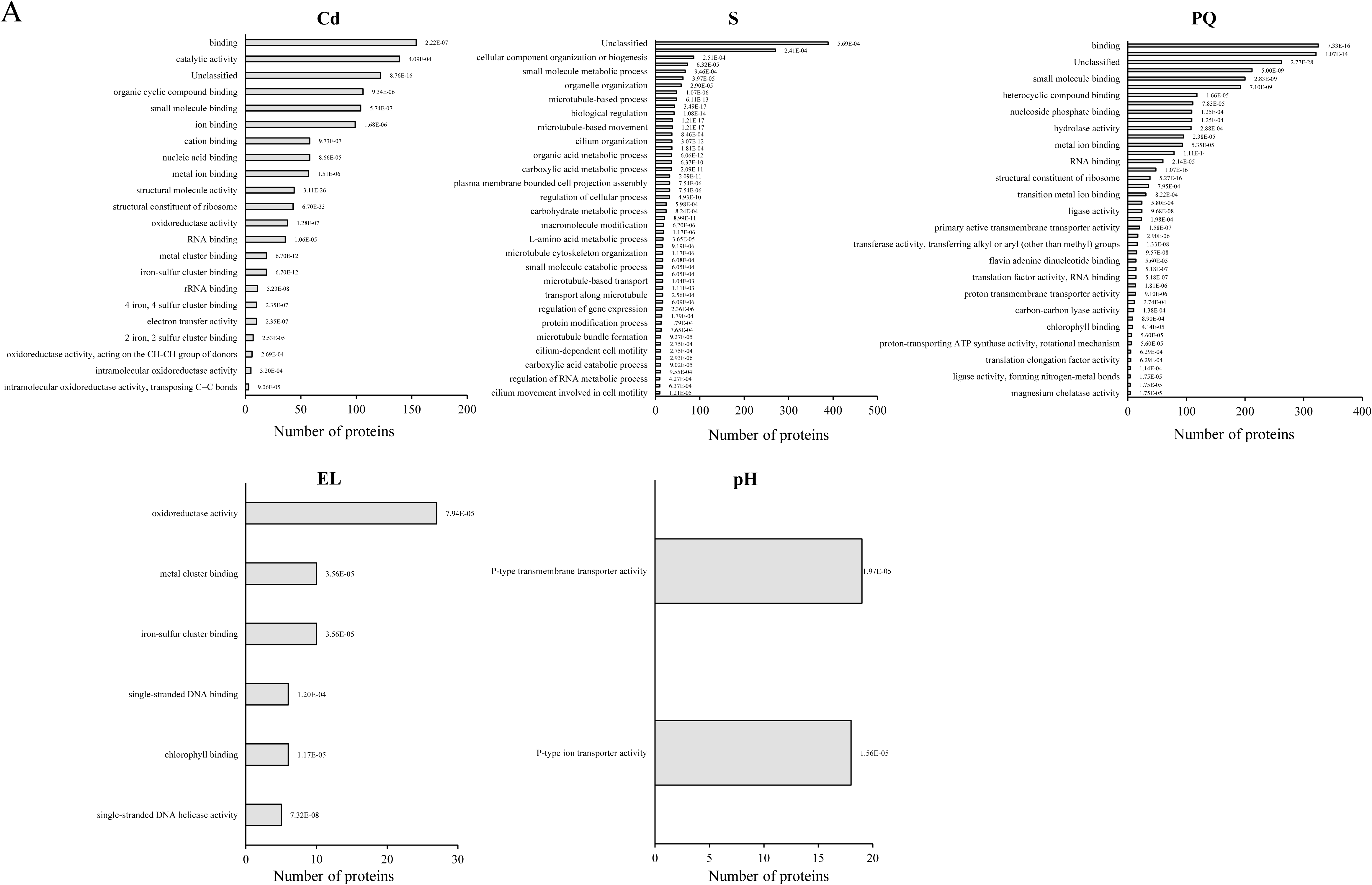
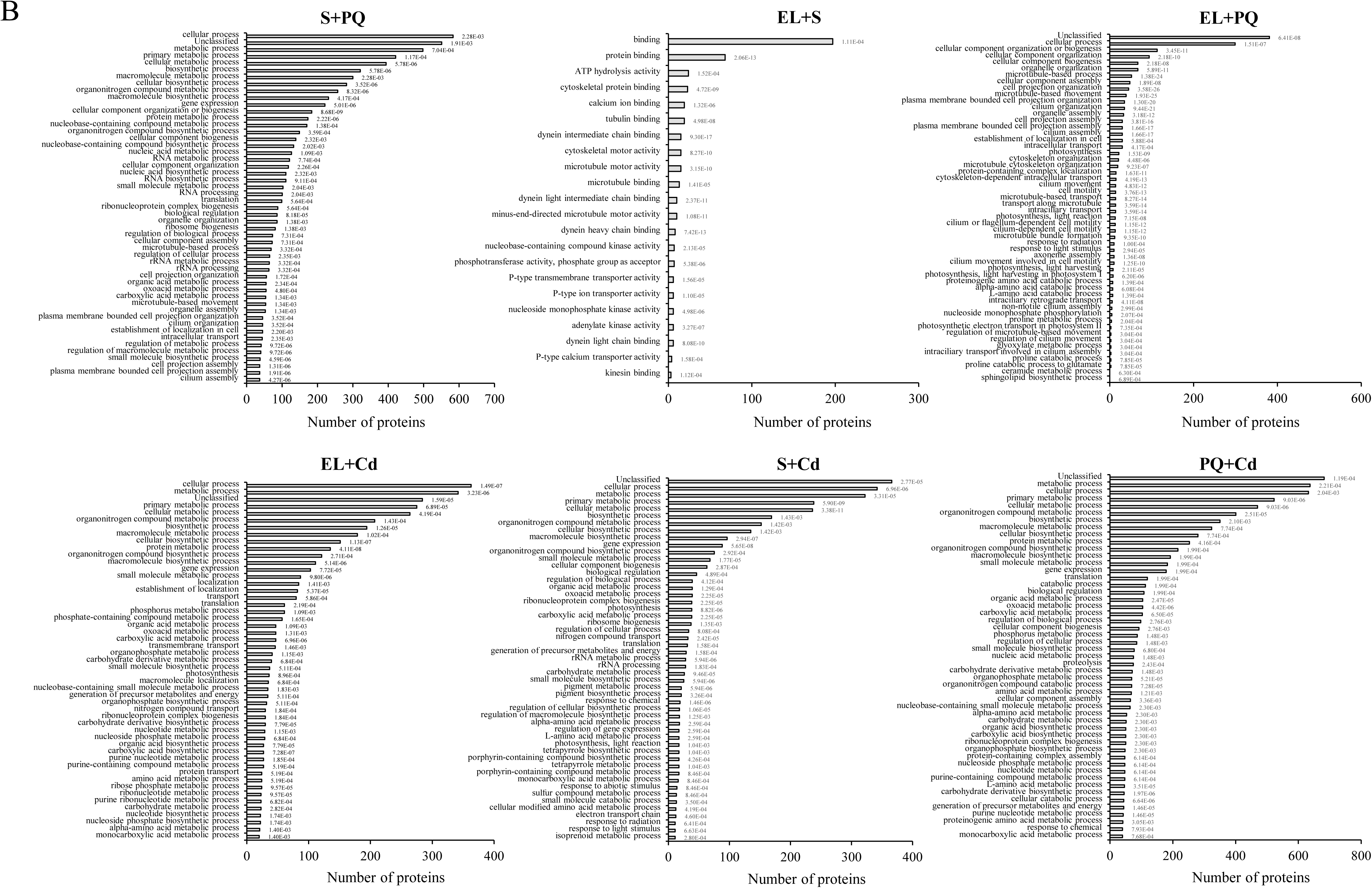
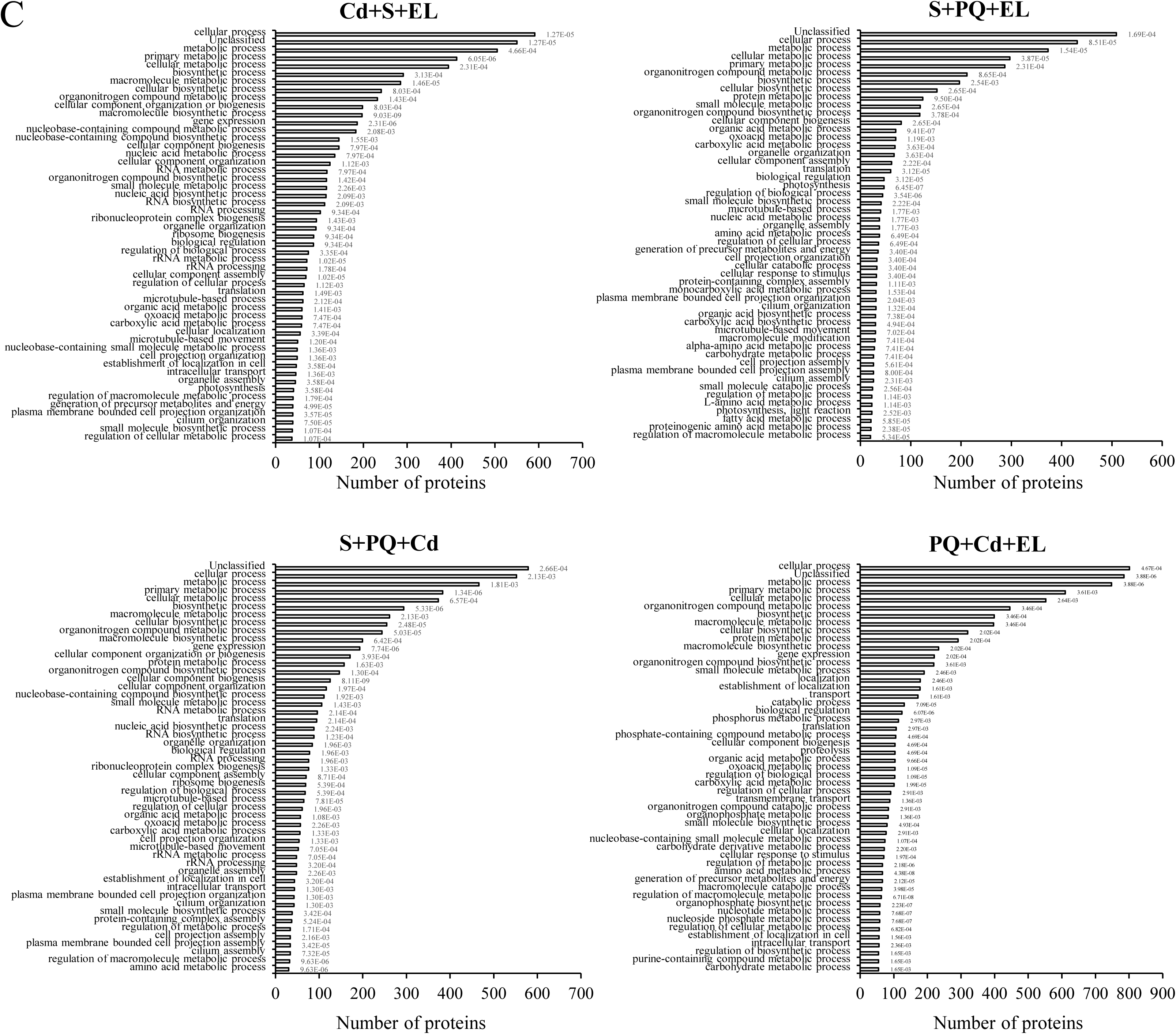
Gene ontology (GO) annotation of respiratory burst oxidase homolog 1 (*rbo1*) *C. reinhardtii* proteins with a significant change in abundance in response to multifactorial stress combination. **A)** GO annotation of proteins with a significant change in abundance in response to treatment with salinity (S), cadmium (Cd), excess light (EL), acidic pH (pH), or paraquat (PQ). **B)** Same as A, but for all 2-stress combinations. **C)** Same as A, but for all 3-, 4-, and 5- stress combinations. In support of Figures 4 and 5. Fold change data is available in the supplemental tables S42 and S43.

**Figure S7.**
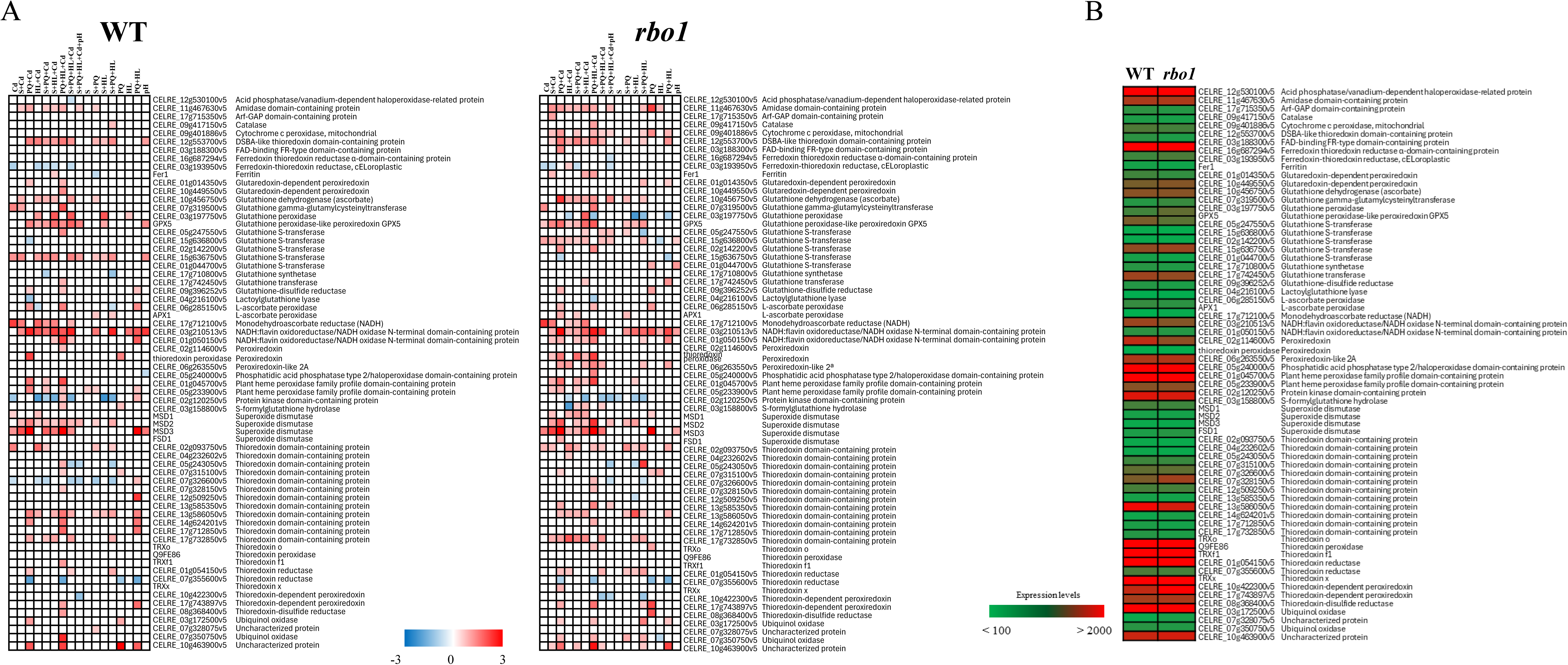
Heatmaps of reactive oxygen species (ROS) metabolizing enzymes abundance in WT and the *rbo1* mutant. **A)** Heatmaps of ROS metabolizing enzymes abundance in WT (left) and *rbo1* (right) *C. reinhardtii* cells subjected to a MFSC of salinity (S), cadmium (Cd), excess light (EL), acidic pH (pH), and/or paraquat (PQ). **B)** Heatmap of protein expression level of the different ROS metabolizing proteins in WT and *rbo1* in the absence of stress (CT). In support of Figure 5.

**Figure S8.**
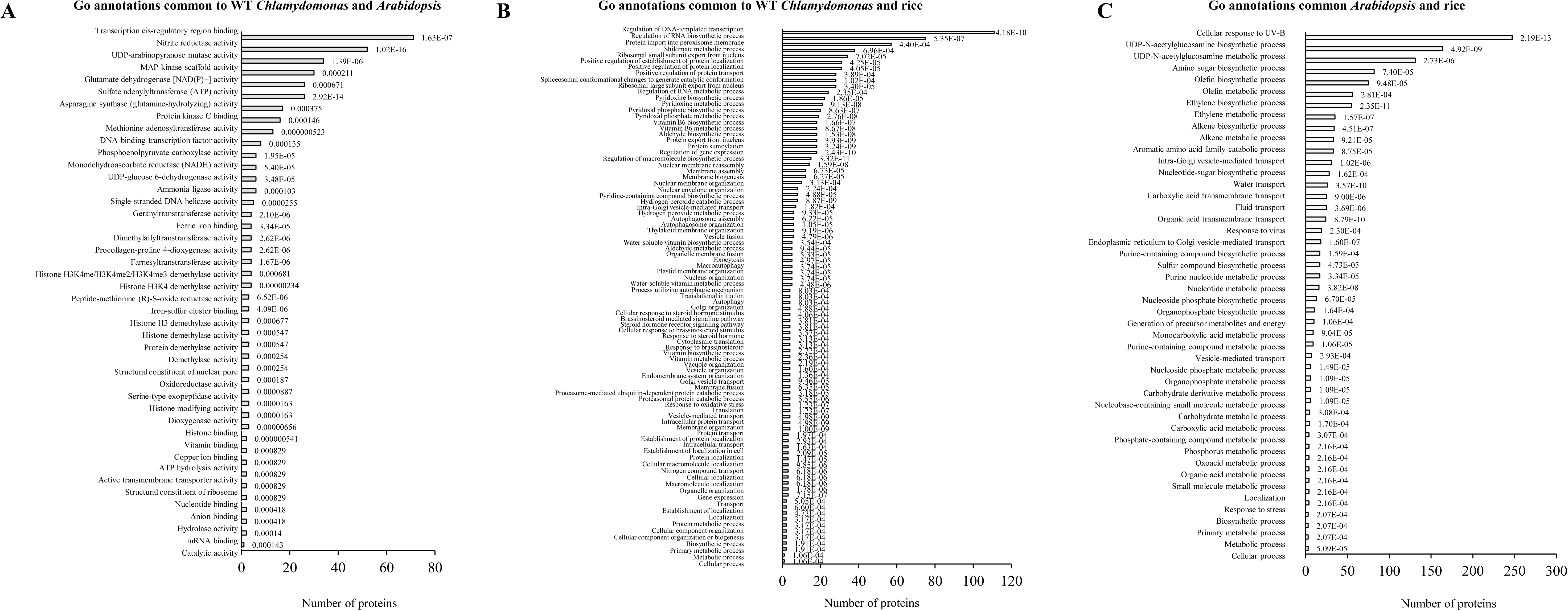
Gene ontology (GO) annotation of conserved proteins with a significant change in abundance in response to multifactorial stress combination. **A)** GO annotation of proteins common to *Arabidopsis* and *C. reinhardtii* from Figure 5D. **B)** GO annotation of proteins common to *C. reinhardtii* and rice from Figure 5D. **C)** GO annotation of proteins common to rice and *Arabidopsis* from Figure 5D. In support of Figure 5.

## List of Supplementary Tables

**Table S1.** Proteins altered in *Chlamydomonas* wild type in response to cadmium (Cd), compared to control.

**Table S2.** Proteins altered in *Chlamydomonas* wild type in response to salinity (S), compared to control.

**Table S3.** Proteins altered in *Chlamydomonas* wild type in response to excess light (EL), compared to control.

**Table S4.** Proteins altered in *Chlamydomonas* wild type in response to paraquat (PQ), compared to control.

**Table S5.** Proteins altered in *Chlamydomonas* wild type in response to pH = 5.5, compared to control.

**Table S6.** Proteins altered in *Chlamydomonas* wild type in response to salinity and paraquat (S+PQ), compared to control.

**Table S7.** Proteins altered in *Chlamydomonas* wild type in response to salinity and cadmium (S+Cd), compared to control.

**Table S8.** Proteins altered in *Chlamydomonas* wild type in response to paraquat and cadmium (PQ+Cd), compared to control.

**Table S9.** Proteins altered in *Chlamydomonas* wild type in response to excess light and salinity (EL+S), compared to control.

**Table S10.** Proteins altered in *Chlamydomonas* wild type in response to excess light and cadmium (EL+Cd), compared to control.

**Table S11.** Proteins altered in *Chlamydomonas* wild type in response to paraquat and excess light (PQ+EL), compared to control.

**Table S12.** Proteins altered in *Chlamydomonas* wild type in response to salinity, paraquat and cadmium (S+PQ+Cd), compared to control.

**Table S13.** Proteins altered in *Chlamydomonas* wild type in response to salinity, cadmium and excess light (S+Cd+EL), compared to control.

**Table S14.** Proteins altered in *Chlamydomonas* wild type in response to salinity, paraquat and excess light (S+PQ+EL), compared to control.

**Table S15.** Proteins altered in *Chlamydomonas* wild type in response to salinity, cadmium and excess light (PQ+Cd+EL), compared to control.

**Table S16.** Proteins altered in *Chlamydomonas* wild type in response to salinity, paraquat, cadmium and excess light (S+PQ+Cd+EL), compared to control.

**Table S17.** Proteins altered in *Chlamydomonas* wild type in response to salinity, paraquat, cadmium, excess light and pH (S+PQ+Cd+EL+pH), compared to control.

**Table S18.** Proteins altered in *Chlamydomonas rbo1* mutant in response to cadmium (Cd), compared to control.

**Table S19.** Proteins altered in *Chlamydomonas rbo1* mutant in response to salinity (S), compared to control.

**Table S20.** Proteins altered in *Chlamydomonas rbo1* mutant in response to excess light (EL), compared to control.

**Table S21.** Proteins altered in *Chlamydomonas rbo1* mutant in response to paraquat (PQ), compared to control.

**Table S22.** Proteins altered in *Chlamydomonas rbo1* mutant in response to pH = 5.5, compared to control.

**Table S23.** Proteins altered in *Chlamydomonas rbo1* mutant in response to salinity and paraquat (S+PQ), compared to control.

**Table S24.** Proteins altered in *Chlamydomonas rbo1* mutant in response to salinity and cadmium (S+Cd), compared to control.

**Table S25.** Proteins altered in *Chlamydomonas rbo1* mutant in response to paraquat and cadmium (PQ+Cd), compared to control.

**Table S26.** Proteins altered in *Chlamydomonas rbo1* mutant in response to excess light and salinity (EL+S), compared to control.

**Table S27.** Proteins altered in *Chlamydomonas rbo1* mutant in response to excess light and cadmium (EL+Cd), compared to control.

**Table S28.** Proteins altered in *Chlamydomonas rbo1* mutant in response to paraquat and excess light (PQ+EL), compared to control.

**Table S29.** Proteins altered in *Chlamydomonas rbo1* mutant in response to salinity, paraquat and cadmium (S+PQ+Cd), compared to control.

**Table S30.** Proteins altered in *Chlamydomonas rbo1* mutant in response to salinity, cadmium and excess light (S+Cd+EL), compared to control.

**Table S31.** Proteins altered in *Chlamydomonas rbo1* mutant in response to salinity, paraquat and excess light (S+PQ+EL), compared to control.

**Table S32.** Proteins altered in *Chlamydomonas rbo1* mutant in response to salinity, cadmium and excess light (PQ+Cd+EL), compared to control.

**Table S33.** Proteins altered in *Chlamydomonas rbo1* mutant in response to salinity, paraquat, cadmium and excess light (S+PQ+Cd+EL), compared to control.

**Table S34.** Proteins altered in *Chlamydomonas rbo1* mutant in response to salinity, paraquat, cadmium, excess light and pH=5.5 (S+PQ+Cd+EL+pH), compared to control.

**Table S35.** Proteins significantly altered in response to 3-, 4- and 5- stress combination in *Chlamydomonas* wild type.

**Table S36.** *Arabidopsis thaliana* homologs of the *Chlamydomonas* proteins significantly altered in response to 3-, 4- and 5- stress combination (from Table S35).

**Table S37.** Significant proteins altered in response to 3-, 4-, and 5- stress combination in rice (data extracted from Sinha et al. 2024).

**Table S38.** Significant transcripts altered in response to 3-, 4-, 5- and 6- multifactorial stress combination in *Arabidopsis thaliana* (data extracted from Zandalinas et al. 2021) and common to significant proteins expressed in WT *Chlamydomonas reinhardtii* in response to 3-, 4- and 5- multifactorial stress combination.

**Table S39.** Significant proteins altered in response to 3-, 4-, and 5- multifactorial stress combination in rice (data extracted from Sinha et al. 2024) and common to significant proteins expressed in WT *Chlamydomonas reinhardtii* in response to 3-, 4- and 5- multifactorial stress combination.

**Table S40.** Significant transcripts altered in response to 3-, 4-, 5- and 6- multifactorial stress combination in *Arabidopsis thaliana* (data extracted from Zandalinas et al. 2021) and common to significant proteins altered in response to 3-, 4-, and 5- multifactorial stress combination in rice (data extracted from Sinha et al. 2024).

**Table S41.** Significant transcripts altered in response to 3-, 4-, 5- and 6- multifactorial stress combination in *Arabidopsis thaliana* (data extracted from Zandalinas et al. 2021) and common to significant proteins altered in response to 3-, 4-, and 5- multifactorial stress combination in rice (data extracted from Sinha et al. 2024) and significant proteins altered in response to 3-, 4-, and 5- multifactorial stress combination in WT *Chlamydomonas*.

**Table S42.** Significant proteins related to ROS wave altered in response to multifactorial stress combination in WT *Chlamydomonas reinhardtii*.

**Table S43.** Significant proteins related to ROS wave altered in response to multifactorial stress combination in *rbo1 Chlamydomonas reinhardtii*.

**Table S44.** Significant proteins related to Fe-S proteins, oxidoreductases, ion channels, photosynthesis proteins and ATPase-coupled transporters altered in response to individual stresses in WT *Chlamydomonas reinhardtii*. Growth rate at 24h is also included.

**Table S45.** Significant proteins related to Fe-S proteins, oxidoreductases, ion channels, photosynthesis proteins and ATPase-coupled transporters altered in response to individual stresses in *rbo1 Chlamydomonas reinhardtii*. Growth rate at 24h is also included.

**Table S46**. Proteins related to ROS wave expression values under non-stress conditions in WT and *rbo1 Chlamydomonas reinhardtii*.

## REFERENCES

1. Altenhoff AM, Train CM, Gilbert KJ, Mediratta I, de Farias TM, Moi D, Nevers Y, Radoykova HS, Rossier V, Vesztrocy AW, et al (2021) OMA orthology in 2021: website overhaul, conserved isoforms, ancestral gene order and more. Nucleic Acids Res 49: D373–D379

2. Anderson A, Laohavisit A, Blaby IK, Bombelli P, Howe CJ, Merchant SS, Davies JM, Smith AG (2016) Exploiting algal NADPH oxidase for biophotovoltaic energy. Plant Biotechnol J 14: 22–28

3. Bailey-Serres J, Parker JE, Ainsworth EA, Oldroyd GED, Schroeder JI (2019) Genetic strategies for improving crop yields. Nature 575: 109–118

4. Bernd KK, Cook N (2002) Microscale assay monitors algal growth characteristics. BioTechniques 32: 1256, 1258–1259

5. de Carpentier F, Lemaire SD, Danon A (2019) When Unity Is Strength: The Strategies Used by *Chlamydomonas* to Survive Environmental Stresses. Cells 8:1307

6. Fichman Y, Rowland L, Oliver MJ, Mittler R (2023) ROS are evolutionary conserved cell-to-cell stress signals. Proc Natl Acad Sci U S A 120: e2305496120

7. Hemker F, Ammelburger N, Jahns P (2024) Intervening dark periods negatively affect the photosynthetic performance of *Chlamydomonas reinhardtii* during growth under fluctuating high light. Plant Cell Environ. 47: 4246–4258.

8. Hoham RW, Remias D (2020) Snow and Glacial Algae: A Review1. J Phycol 56: 264–282

9. Intergovernmental Panel on Climate Change (IPCC) (2023) Climate Change 2022 – Impacts, Adaptation and Vulnerability: Working Group II Contribution to the Sixth Assessment Report of the Intergovernmental Panel on Climate Change. Clim Chang 2022 – Impacts, Adapt Vulnerability. doi: 10.1017/9781009325844

10. Lesk C, Rowhani P, Ramankutty N (2016) Influence of extreme weather disasters on global crop production. Nat 529: 84–87

11. Li J, Mu J, Bai J, Fu F, Zou T, An F, Zhang J, Jing H, Wang Q, Li Z, et al (2013) PARAQUAT RESISTANT1, a Golgi-Localized Putative Transporter Protein, Is Involved in Intracellular Transport of Paraquat. Plant Physiol 162: 470–483

12. Liu K, Ding X, Wang HF, Zhang X, Hozzein WN, Wadaan MAM, Lan A, Zhang B, Li W (2014) Eukaryotic microbial communities in hypersaline soils and sediments from the alkaline hypersaline Huama Lake as revealed by 454 pyrosequencing. Antonie Van Leeuwenhoek 105: 871–880

13. Ma X, Wei H, Zhang Y, Duan Y, Zhang W, Cheng Y, Xia XQ, Shi M (2020) Glutathione peroxidase 5 deficiency induces lipid metabolism regulated by reactive oxygen species in *Chlamydomonas reinhardtii*. Microb Pathog. 147:104358.

14. Márquez-Zacarías P, Conlin PL, Tong K, Pentz JT, Ratcliff WC (2021) Why have aggregative multicellular organisms stayed simple? Curr Genet 67: 871–876

15. Mittler R (2006) Abiotic stress, the field environment and stress combination. Trends Plant Sci 11: 15–19

16. Mittler R, Zandalinas SI, Fichman Y, Van Breusegem F (2022) Reactive oxygen species signalling in plant stress responses. Nat Rev Mol Cell Biol 23: 663–679

17. Pascual LS, Mittler R, Sinha R, Peláez-Vico MÁ, López-Climent MF, Vives-Peris V, Gómez-Cadenas A, Zandalinas SI (2023) Jasmonic acid is required for tomato acclimation to multifactorial stress combination. Environ Exp Bot 213: 105425

18. Peláez-Vico MÁ, Sinha R, Induri SP, Lyu Z, Venigalla SD, Vasireddy D, Singh P, Immadi MS, Pascual LS, Shostak B, et al (2023) The impact of multifactorial stress combination on reproductive tissues and grain yield of a crop plant. Plant J 117:1728–1745

19. Ratcliff WC, Herron MD, Howell K, Pentz JT, Rosenzweig F, Travisano M (2013) Experimental evolution of an alternating uni- and multicellular life cycle in *Chlamydomonas reinhardtii*. Nat Commun 4: 1–7

20. Rhee SY, Anstett D, Cahoon E, Covarrubias-Robles AA, Danquah E, Doudareva N, Ezura H, Gilbert KJ, Gutiérrez RA, Heck M, et al (2024) Resilient plants, sustainable future. Trends Plant Sci In press

21. Richardson K, Steffen W, Lucht W, Bendtsen J, Cornell SE, Donges JF, Drüke M, Fetzer I, Bala G, von Bloh W, et al (2023) Earth beyond six of nine planetary boundaries. Sci Adv 9:eadh2458

22. Rillig MC, Ryo M, Lehmann A (2021) Classifying human influences on terrestrial ecosystems. Glob Chang Biol 27: 2273–2278

23. Rillig MC, Ryo M, Lehmann A, Aguilar-Trigueros CA, Buchert S, Wulf A, Iwasaki A, Roy J, Yang G (2019) The role of multiple global change factors in driving soil functions and microbial biodiversity. Science 366: 886–890

24. Sage RF (2020) Global change biology: A primer. Glob Chang Biol 26: 3–30

25. Sasso S, Stibor H, Mittag M, Grossman AR (2018) From molecular manipulation of domesticated *Chlamydomonas reinhardtii* to survival in nature. Elife 7: e39233

26. Sinha R, Peláez-Vico MÁ, Shostak B, Nguyen TT, Pascual LS, Ogden AM, Lyu Z, Zandalinas SI, Joshi T, Fritschi FB, et al (2024) The effects of multifactorial stress combination on rice and maize. Plant Physiol 194: 1358–1369

27. Smirnoff N, Arnaud D (2019) Hydrogen peroxide metabolism and functions in plants. New Phytol 221: 1197–1214.

28. Speißer B, Wilschut RA, van Kleunen M (2022) Number of simultaneously acting global change factors affects composition, diversity and productivity of grassland plant communities. Nat Commun 13:7811

29. Tong K, Bozdag GO, Ratcliff WC (2022) Selective drivers of simple multicellularity. Curr Opin Microbiol 67: 102141

30. Youssef WA, Feil R, Saint-Sorny M, Johnson X, Lunn JE, Grimm B, Brzezowski P (2023) Singlet oxygen-induced signalling depends on the metabolic status of the *Chlamydomonas reinhardtii* cell. Commun Biol. 6:529.

31. Zandalinas SI, Mittler R (2022) Plant responses to multifactorial stress combination. New Phytol 234: 1161–1167

32. Zandalinas SI, Sengupta S, Fritschi FB, Azad RK, Nechushtai R, Mittler R (2021) The impact of multifactorial stress combination on plant growth and survival. New Phytol 230: 1034–1048

33. Zhang H, Zhu J, Gong Z, Zhu JK (2021) Abiotic stress responses in plants. Nat Rev Genet 23: 104–119

34. Zhang N, Mattoon EM, McHargue W, Venn B, Zimmer D, Pecani K, Jeong J, Anderson CM, Chen C, Berry JC, et al (2022) Systems-wide analysis revealed shared and unique responses to moderate and acute high temperatures in the green alga *Chlamydomonas reinhardtii*. Commun Biol 5: 5(1):460

35. Zhou Y, Fichman Y, Zhang S, Mittler R, Chen SJ (2024) Modeling the reactive oxygen species (ROS) wave in Chlamydomonas reinhardtii colonies. Free Radic Biol Med 222: 165–172

